# An Effector Index to Predict Causal Genes at GWAS Loci

**DOI:** 10.1101/2020.06.28.171561

**Authors:** Vincenzo Forgetta, Lai Jiang, Nicholas A. Vulpescu, Megan S. Hogan, Siyuan Chen, John A. Morris, Stepan Grinek, Christian Benner, Dong-Keun Jang, Quy Hoang, Noel Burtt, Jason A. Flannick, Mark I McCarthy, Eric Fauman, Celia MT Greenwood, Matthew T. Maurano, J. Brent Richards

**Affiliations:** Centre for Clinical Epidemiology, Lady Davis Institute for Medical Research, Jewish General Hospital, Montreal, Canada; Departments of Medicine, Epidemiology and Biostatistics, McGill University, Montreal, Canada; Institute for Systems Genetics and Department of Pathology, NYU School of Medicine, New York, USA; New York Genome Center, New York, USA; Department of Biology, New York University, New York, USA; Institute for Molecular Medicine Finland (FIMM), University of Helsinki, 00014 Helsinki, Finland; Program in Medical and Population Genetics, Metabolism Program, Broad Institute of Harvard and MIT, Cambridge, MA, USA; Department of Pediatrics, Harvard Medical School, Boston, MA; Division of Genetics and Genomics, Boston Children’s Hospital, Boston, MA; Wellcome Centre for Human Genetics, University of Oxford, UK; Internal Medicine Research Unit, Pfizer Worldwide Research, Development and Medical, USA; Gerald Bronfman Department of Oncology, McGill University, Montréal, Canada; Department of Human Genetics, McGill University, Montréal, Canada; Department of Medicine, McGill University, Montréal, Canada; Department of Twin Research, King’s College London, London. United Kingdom

## Abstract

Drug development and biological discovery require effective strategies to map existing genetic associations to causal genes. To approach this problem, we began by identifying a set of positive control genes for 12 common diseases and traits that cause a Mendelian form of the disease or are the target of a medicine used for disease treatment. We then identified a widely-available set of genomic features enriching GWAS-associated single nucleotide variants (SNVs) for these positive control genes. Using these features, we trained and validated the Effector Index (*Ei*), a causal gene mapping algorithm using the 12 common diseases and traits. The area under *Ei’s* receiver operator curve to identify positive control genes was 80% and area under the precision recall curve was 29%. Using an enlarged set of independently curated positive control genes for type 2 diabetes which included genes identified by large-scale exome sequencing, these areas increased to 85% and 61%, respectively. The best predictors were coding or transcript altering SNVs, distance to gene and open chromatin-based metrics. We provide the *Ei* algorithm for its widespread use and have created a web-portal to facilitate understanding of results. This work outlines a simple, understandable approach to prioritize genes at GWAS loci for functional follow-up and drug development.

**Author summary:** In order to derive biological insight, or develop drugs based on genome-wide association studies (GWAS) data, causal genes at associated loci need to be identified. GWAS usually identify large genome regions containing many genes, but seldomly identifies specific causal genes. We have developed an algorithm to predict which genes in a region of disease association are likely causal and have named this algorithm the Effector Index. The Effector Index was optimized on diseases that have known causal or drug target genes, and further validated to predict these types of genes in independent datasets. The Effector Index formalizes these predictive features into a tool that can be used by researchers, and results from the traits and diseases studied here are available via the Accelerating Medicine Partnership web-portal at http://hugeamp.org/effectorgenes.html.

## Introduction

The majority of late-stage drug development programs fail.[1–3] The most common cause of these failures is a lack of efficacy of the medicine on the disease outcome.[2] Such failures are due, in part, to unreliable drug target identification and validation.[4] Recent evidence has suggested that when drug development programs have support from human genetics the probability of success increases.[5–7] Yet, this evidence relies upon retrospective surveys of drug development programs and assumes that human genetic associations, which are most often non-coding, could be used to identify the specific causal gene(s) at an associated locus—while in reality this is challenging. Moving forward, the use of human genetics for target identification and validation will require that a genetic association be mapped to specific genes. Therefore, strategies to reliably map the thousands of GWAS associations to causal genes are required to realize the potential of human genetics to deliver medicines to the clinic.

Many strategies have been developed to prioritize genes at GWAS loci[8–11] each leveraging one or more biological data sets under specific assumptions (**Table 1**), primarily by using SNVs that overlap, or are near, specific types of annotations that have been shown to be enriched for proximity to genes of known biological relevance to the trait[12,13]. These methods excel at their intended purposes: to identify genes that may be influenced by eQTLs, to identify genes with known biological purposes, or to identify genes near genomic annotations of interest. However, bulk tissue *cis*-eQTLs account for a small percentage of disease heritability and genes with large effects on complex traits tend to have low *cis* heritability[14]. While the other methods rely upon known biology, thereby limiting the ability to discover new pathways and drug targets. Most importantly, these methods have not been evaluated against a set of positive control genes most likely to be relevant for drug development, which would contain the targets of drugs successfully used in the clinic and genes that cause Mendelian forms of the complex disease.

**Table 1.** High-level conceptual summary of approaches for gene prioritization at GWAS loci. Shown are main classes of methods, exemplified by a published implementation.

| Class | Example Method | Annotation Data Source | Key Assumption | Key Goal |
| --- | --- | --- | --- | --- |
| eQTL | eCAVIAR[8] | eQTL | Target genes demonstrate heritable expression differences in eQTL tissues | Colocalize GWAS and eQTL signals to detect target genes |
| Guilt-by-association | DEPICT[9] | Gene-centric annotation (e.g. curated gene sets or gene expression sets) | Multiple associations will be driven by genes sharing functional annotation | Identify gene sets demonstrating functional enrichment across GWAS loci |
| Gene set enrichment analysis | MAGENTA[10] | Curated gene sets (KEGG, REACTOME) | Causal genes can be prioritized by known biological annotations |  |
| Functional priors in statistical fine mapping | PAINTOR[11] | Statistical fine-mapping, transcript, and functional genomics data | Genes near credible SNVs with functional impact are likely causal | Prioritize genetic variants using statistical finemapping and functional annotation. |
| Integration of functional genomics data using supervised learning | Effector Index (Ei) | Statistical fine-mapping, transcript, eQTL, DHS data | Building on key assumptions of other tools, find specific patterns in the annotations or distance of credible SNVs around genes can predict their probability of being causal | Using a curated causal gene set of drug targets or Mendelian disease genes, train a complex statistical model to predict causal genes that jointly considers the functional annotation and distance of SNVs at the locus. |

An idealized strategy to map genetic associations to causal genes would have several characteristics. First, the predictive features that enable mapping a genetic association to a causal gene would be easily and reliably measured, shared across diseases and available from public datasets. By using a parsimonious set of simply calculated predictors, such a mapping tool would be more easily applicable. Second, the strategy would be generalizable across diseases, enabling its use even in diseases where few causal genes are known. Third, the strategy would be evaluated by testing its predictive ability against a set of truly causal positive control genes. Without such a set of causal positive control genes, the value of any strategy cannot be properly examined. Last, the strategy would be evaluated in a dataset distinct from the training dataset. We therefore developed the “Effector Index (*Ei*)”, an algorithm which generates the probability of causality for all genes at a GWAS locus. Specifically, the *Ei* aims to answer the question, “What is the probability of causality for each gene at a locus which harbors genome-wide significant SNVs for a disease or trait?”.

The *Ei* provides a rapid and readily understandable way to assess the probability that a gene at a GWAS locus is causal for a disease or trait. It requires relatively simple annotations from GWAS summary statistics, genomic positions and DNaseI hypersensitive sites (DHS). These findings may accelerate drug development by providing guidance on which gene(s) at a GWAS locus are likely to be causal, and they should also prioritize genes for downstream functional genomic and biological experiments. Below we present the key results of the study.

## Results

### Overall Strategy

To generate the *Ei*, we first carefully defined a set of positive control causal genes for 12 diseases and traits by relying only upon data from humans, defining positive control genes as genes whose perturbation causes a Mendelian form of the common disease, or whose encoded protein acts as a drug target for the common disease. We next assessed the genomic annotations at GWAS loci that enriched for positive control genes. Then, to predict each gene’s probability of causality at a GWAS locus we used both locus-level features, such as the number of genes at a locus, and gene-level features, such as distance of a gene to nearest associated SNV. We trained a gradient boosted trees algorithm, as implemented in XGBoost[15] to generate the probability of causality for each gene at each GWAS locus for 11 diseases and tested the resulting model on the 12^th^ disease, iterating this process across all 12 diseases. As a sensitivity analysis, for type 2 diabetes (T2D) we tested an enlarged a set of positive control genes that, in addition to the positive controls as described above, included independently and manually curated positive controls, incorporating recent evidence from coding variants arising from exome array and exome sequencing studies[16] (http://www.type2diabetesgenetics.org/).

### Diseases and Traits Studied, and Positive Control Genes

We selected a broad panel of diseases and quantitative traits which have been the subject of large-scale GWASs, such as those from UK Biobank or large-scale international GWAS consortia[13] (**Table 2**). The diseases and traits studied included: T2D, low density lipoprotein (LDL) cholesterol level, adult height, calcium level, hypothyroidism, triglyceride (Tg) level, estimated bone mineral density (eBMD), glucose level, red blood cell count (RBC) systolic blood pressure (SBP), diastolic blood pressure (DBP) and direct bilirubin level.

**Table 2.** Summary of GWAS studies.

| Phenotype | Data Source | N<br>individuals | Harmonized<br>SNVs* | Lead<br>SNPs‡ | Loci§ | Finemapped<br>SNVs¶ | Positive<br>Control<br>Genes |  |
| --- | --- | --- | --- | --- | --- | --- | --- | --- |
|  |  |  |  |  |  |  | OMIM | Drug |
| Type 2 diabetes | Mahajan et al. (2018) | 898,130 | 9,564,286 | 226 | 147 | 679 | 48 | 7 |
| Estimated bone mineral density | Morris et al. (2018) | 426,824 | 8,540,200 | 2,170 | 700 | 5,514 | 54 | 14 |
| Diastolic blood pressure | Price lab, UK Biobank† | 376,437 | 8,812,132 | 790 | 460 | 2,127 | 15 | 18 |
| Height | Price lab, UK Biobank† | 408,092 | 8,811,966 | 4,155 | 1,011 | 11,698 | 65 | 2 |
| Hypothyroidism | Price lab, UK Biobank† | 459,324 | 8,811,971 | 166 | 125 | 461 | 7 | 6 |
| Red blood cell count | Price lab, UK Biobank† | 445,174 | 8,811,899 | 1,310 | 571 | 3,021 | 66 | 6 |
| Systolic blood pressure | Price lab, UK Biobank† | 376,437 | 8,812,132 | 812 | 481 | 2,269 | 15 | 18 |
| Calcium | UK Biobank, this study | 380,228 | 10,545,875 | 438 | 251 | 1,976 | 4 | 11 |
| Direct bilirubin | UK Biobank, this study | 349,743 | 10,545,843 | 222 | 78 | 561 | 2 | 0 |
| Glucose | UK Biobank, this study | 375,396 | 10,545,731 | 147 | 106 | 568 | 71 | 20 |
| Low density lipoprotein | UK Biobank, this study | 333,541 | 10,223,520 | 586 | 243 | 2,201 | 17 | 5 |
| Triglycerides | UK Biobank, this study | 333,891 | 10,223,638 | 454 | 224 | 1,587 | 17 | 6 |

\*Count for SNPs present in UK Biobank and with MAF > 0.005.
†https://data.broadinstitute.org/alkesgroup/UKBB/UKBB_409K/
‡LD independent SNPs at p-value < $5 \times 10^{-8}$ and $r^2 < 0.01$ , excluding SNPs within the MHC locus and those from non-converged FINEMAP loci
§Merged loci containing one or more lead SNPs within 50 Kbp of each other. Excludes loci that overlap the MHC and those that failed convergence during FINEMAP.
¶FINEMAP SNVs with $\log_{10}(\text{BF}) >$
OMIM: Count of genes identified through the Online Mendelian Inheritance in Man to cause a Mendelian form of the disease, or influence the trait
Drug: Count of known drug targets used to treat each disease, or influence the trait

We required that for each disease or trait, a set of stringently defined “positive control genes” could be identified meeting at least one of two criteria: (i) perturbation of the gene is known to cause of a Mendelian form of the disease (or influences the trait); or (ii) the gene’s protein is the target of a therapy successfully developed to treat the disease, or influence the trait. To identify Mendelian disease genes, we first used the Human Disease Ontology[17] database to identify diseases influencing traits studied (**Table S1**). Using the resulting list of curated ontological terms we obtained a list of Mendelian disease genes from the Online Mendelian Inheritance in Man (OMIM; **Table S2**).[18] Positive control drug targets were identified by first collecting guideline-suggested medications for each trait or disease, as described in UpToDate, an online decision tool written and edited by medical experts (**Table S3**),[19] then we gathered the known targets of these medicines using DrugBank (**Tables S4**).[20]

We identified 494 positive control genes across the 12 diseases and traits, 381 known to cause Mendelian forms of the disease (or influence the trait) and 113 drug targets (**Tables S2 & S4**). This ranged from two for direct bilirubin to 66 for RBC, with an average of 32 per disease/trait. Of the 113 positive control genes from drug targets, we found an average of 9 drug classes per disease trait. This represented 55 unique drug targets, since different classes of clinically used medicines may have the same drug target (**Table S4**).

### Fine-mapping GWAS loci

We applied a statistical fine mapping pipeline to prioritize SNVs from previously published GWASs for T2D,[13] eBMD[12] and *de-novo* GWASs for the other diseases and traits from the UK Biobank (**Fig. 1a**) (**Table 2**). Fine-mapping is more helpful in this context than conditional analyses since it provides probabilistic measures of causality for SNVs.[21] SNVs passing quality control and having a minor allele frequency > 0.005 were retained for LD clumping, followed by merging with adjacent association signals (see **Materials and Methods** for further details). We applied the program FINEMAP[21] using a matching LD panel comprised of 50,000 individuals of white-British ancestry from UK Biobank. These GWASs yielded between 78 and 1,011 independent loci where the average was 366 (**Table 2**). Therefore, each disease or trait had a large number of loci which could be used to train or test the predictive model.

**Fig. 1.**
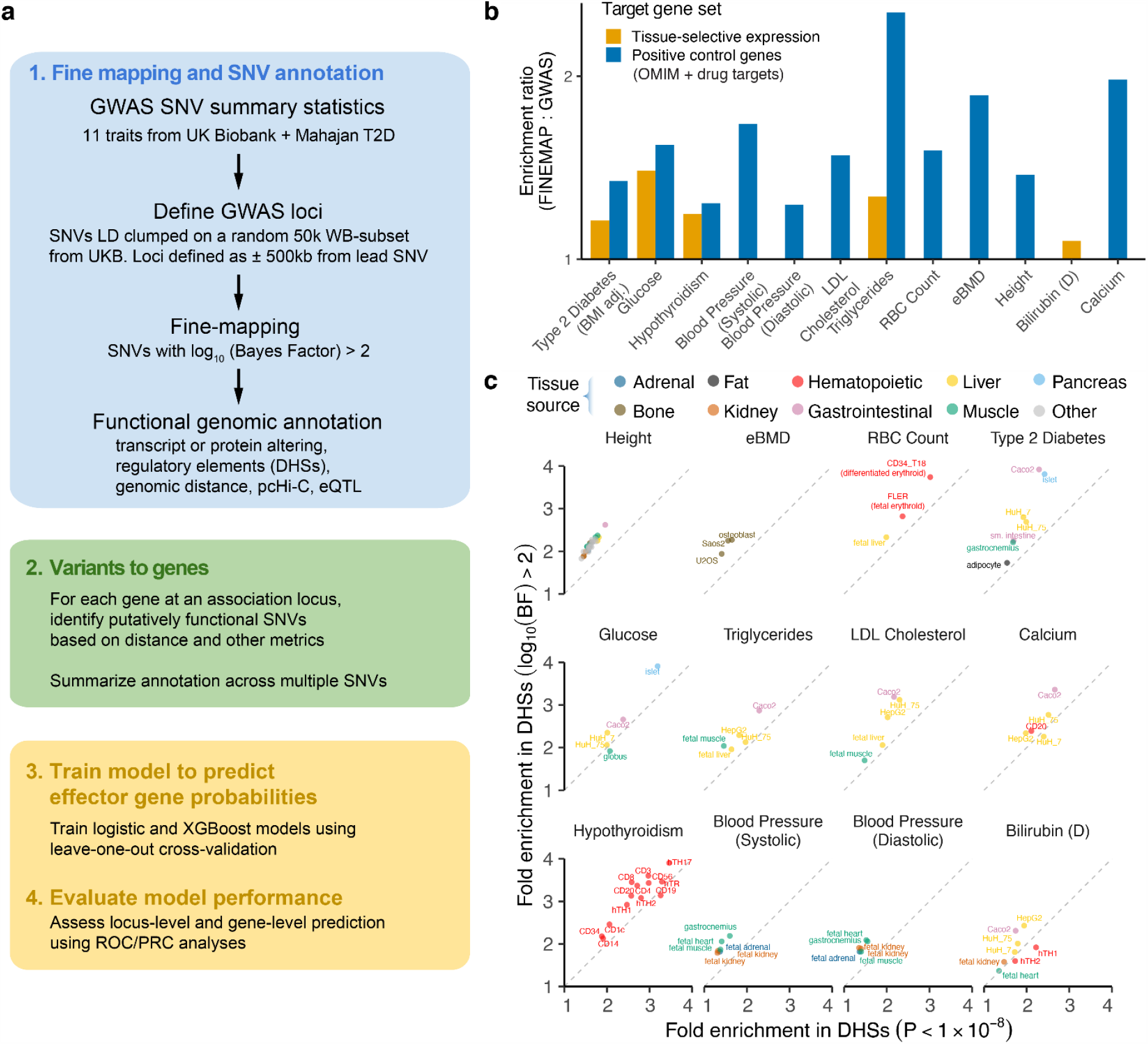
Building the Effector Index and enrichment for likely target genes by statistical fine-mapping. (**a**) Flow diagram depicting: (1) how data was generated using fine-mapping of GWAS summary statistics, followed by SNV annotation and pairing to genes at each GWAS locus (2-3) how this data is used to generate gene- and locus-level features, followed by fitting their feature weights within the models using a leave-one-out analysis, and (4) assessing the performance of the models to predict target genes for loci containing positive control genes. (**b**) Ratio of enrichment for positive control genes within ±25 Kbp of genome-wide significant SNVs (P < 5 × 10^−8^) compared to SNVs having log_10_(BF) > 2 after fine-mapping. Fold enrichment was calculated as the proportion of positive control genes targeted to the proportion of all genes targeted. (**c**) Comparison of enrichments for genome-wide significant SNVs (x-axis) vs. SNVs with log_10_(BF) > 2 SNVs (y-axis) within trait-specific DHS sites (see **Materials and Methods**).

Fine-mapping dramatically reduced the size of the putatively causal SNV set relative to the initial GWAS (**Supplementary Fig. 1a**). Fully 48% of most strongly associated lead SNVs also had the highest FINEMAP log_10_(Bayes Factor (BF)) at a locus, and an average of 55% of these lead SNVs were also in the fine-mapped credible set (**Supplementary Fig. 1b-c**). There was an average of 3.7 fine-mapped SNVs per locus (**Supplementary Fig. 2**), suggesting a large degree of allelic heterogeneity. Consequently, this fine-mapping step substantially reduced the number of SNVs to be considered for mapping to causal genes.

### Statistical fine mapping strongly enriches for positive control genes

We assessed the utility of fine-mapping for enrichment for positive control genes by comparing enrichment using our approach to previously published studies for eBMD[12] and T2D[13]. SNVs achieving a log_10_(BF) > 2 upon fine-mapping demonstrated increased enrichment relative to published credible sets for our positive control gene sets (**Supplementary Fig. 3**). We next compared the enrichment of SNVs for proximity to positive control genes (within 25 Kbp) when SNVs were limited to only those which were genome-wide significant (P < 5 × 10^−8^), compared to fine-mapped SNVs. In every case, the fine-mapped SNVs showed higher enrichment for positive control genes (**Fig. 1b, Supplementary Fig. 4**) compared to genome-wide significant SNVs. Taken together, these findings suggest that the fine-mapping approach decreases the total number of SNVs to be mapped to genes and helps to identify positive control genes. Assuming that each GWAS locus reflects a causal biological signal, these findings indicate that most of the causal genes at GWAS loci are not currently known to cause Mendelian forms of the disease, or act as drug targets, thereby providing the opportunity to identify novel causal genes.

### Assessment of gene expression as a source of positive control genes

Tissue-specific expression has also been used as a method to identify putatively causal genes at GWAS loci,[22,23] and we tested this in a sensitivity analysis as an alternative source for positive control genes. To do so we developed tissue-selective gene sets based on expression using RNA-seq data for a variety of tissues from the GTEx project and purified hematopoietic cells from the ENCODE project (**Supplementary Fig. 5a, Materials and Methods**). We similarly observed that fine-mapped SNVs enriched for tissue-specific expression gene sets (**Supplementary Fig. 5b**), when compared to SNVs surpassing a P-value threshold < 5 × 10^−8^. However, fold-enrichment was substantially higher for the set of positive control genes derived from Mendelian forms of disease and drug targets, than gene sets identified through tissue-selective gene expression (**Supplementary Fig. 4**). Specifically, enrichment of log_10_(BF) > 2 SNVs for the positive control genes identified using Mendelian disease and drug targets was, on average, 6.8-fold higher than enrichment for the expression-derived genes (**Supplementary Fig. 4**), but the tissue-specific expression sets contained 5.4-142 times more genes. Given the clearly stronger enrichment for positive control genes, we did not further consider genes identified through tissue-selective expression as positive control genes.

### Fine-mapping enriches for cell-type selective DNase I hypersensitive sites

It has been shown previously that trait-associated variants localize to genomic regulatory regions of relevant cell and tissue types[24]. To use this enrichment effect as an orthogonal method to validate assumptions inherent to fine-mapping, we analyzed potential local regulatory effects of non-coding disease and trait-associated SNVs by comparison with DNase-seq data from a broad set of cell and tissue types generated the ENCODE and Roadmap Epigenomics projects.[25,26] We also generated DNase-seq data for Saos-2 and U2OS osteosarcoma cell lines and downloaded published accessible sites for pancreatic islets.[27] All data were analyzed using a uniform mapping and peak-calling algorithm (**Materials and Methods**).

We then calculated the enrichment for disease-and trait-associated SNVs in DHSs for each cell or tissue type (**Supplementary Fig. 6**) for progressively increasing log_10_(BF) thresholds. We identified strong enrichment at higher log_10_(BF) cutoffs for DHSs from cell types relevant to the trait. We then selected cell and tissue types which showed strong enrichment for each trait (**Table S5**). By comparing enrichment in DHSs for SNVs with P-value < 1 ×10^−8^ or log_10_(BF) > 2, enrichment was an average of 2-fold higher after fine-mapping (**Fig. 1c)**. These results provide additional orthogonal evidence for the value of statistical fine-mapping in order to map genetic associations to positive control genes.

### Genomic landscape annotations that enrich for positive control genes at GWAS loci

Fine-mapped SNVs were mapped to genes based on several different types of genomic annotations: (i) when the SNV alters amino acid sequence or transcript structure, (ii) overlap of the SNV with DHSs in relevant cell or tissue types, (iii) overlap of the SNV with an eQTL in a relevant tissue, and (iv) promoter-capture Hi-C in a relevant tissue. Most positive control genes in the proximity of fine-mapped SNVs were not targeted directly by a protein or transcript-altering SNV (**Fig. 2a**). But while a sizable proportion of positive control genes were only found by looking at longer range or with chromatin interaction data (**Fig. 2a**), enrichment decreased with distance to TSS (**Fig. 2b**). High-resolution examination of enrichment showed a strong inflection point at close range to the TSS (<50 Kbp); however, enrichment remained substantial even at distances >250 Kbp (**Fig. 2c, Supplementary Fig. 7**). These characteristics became even clearer upon fine-mapping and subsequent restriction to SNVs in cell-type specific DHSs. Enrichment for non-coding SNVs in DHSs was highest at medium range (<25 Kbp away) (**Fig. 2c**). However, while enrichment for more distant SNVs was lower, 69% of positive control genes at GWAS loci were > 25 Kbp from the nearest log_10_(BF) > 2 SNV (**Fig. 2a**). eQTL and promoter capture Hi-C data showed enrichment even after accounting for distance to TSS (**Supplementary Fig. 8**), but the overall magnitude of enrichment was considerably lower than when using simpler distance to gene metrics. While the these genomic features have been previously shown to be enriched at GWAS loci[13], it remains unclear how to systematically weigh their relevance across loci for different traits and diseases. In the following section we demonstrate a model integrating these and other annotations to predict causal genes.

**Fig. 2.**
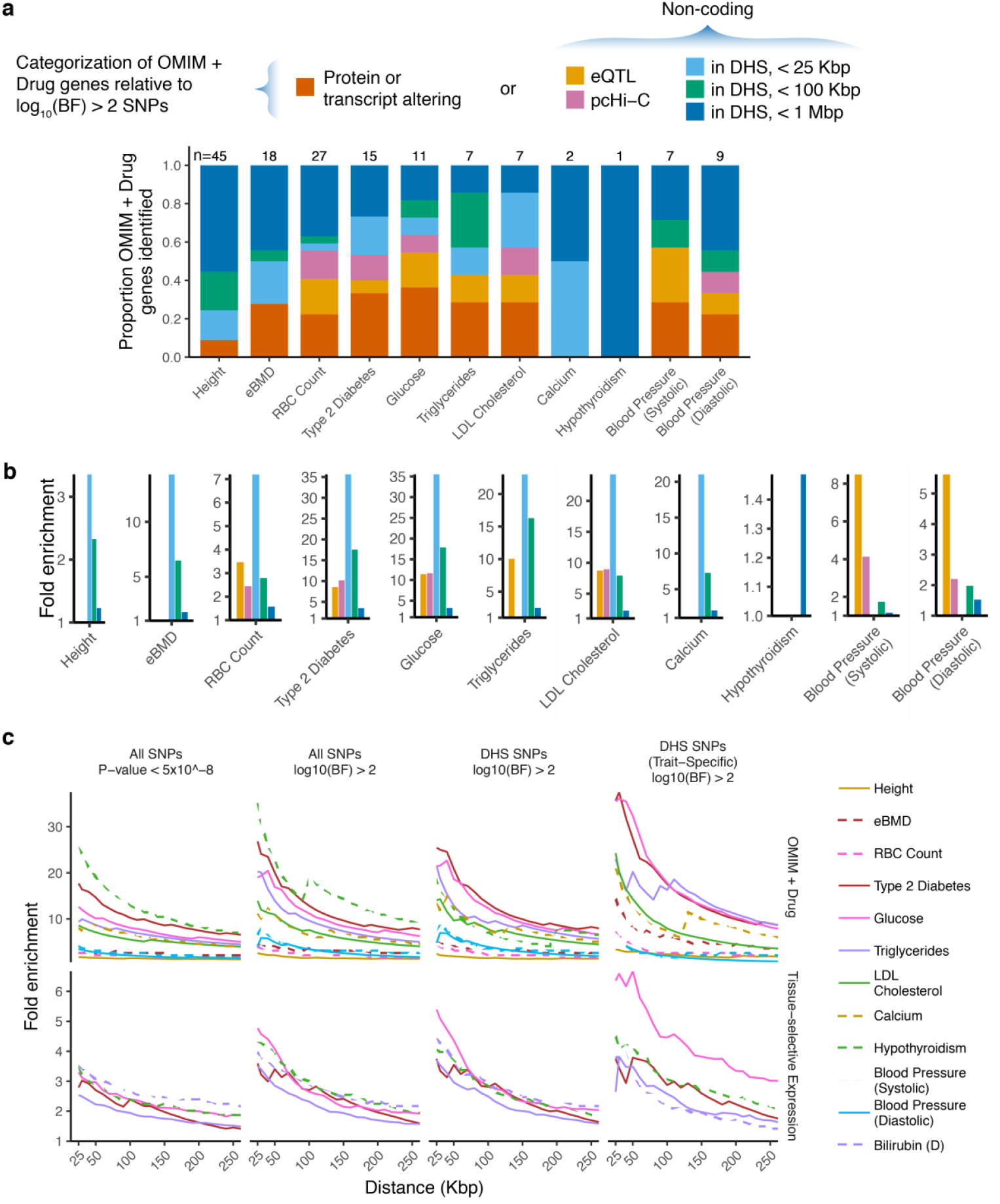
Enrichment of genomic landscape features with positive control genes. Genes with protein or transcript altering SNVs were assessed separately. Non-coding SNVs were classified by overlap with trait-specific DHSs, distance to the TSS, and eQTL or pcHi-C evidence. (**a**) Summary of positive control genes at GWAS loci by relation to log_10_(BF) > 2 SNVs. Bar charts demonstrate the proportion of positive control genes identified by intersection of fine-mapped SNVs with different genomic landscape features. Results are separated by trait/disease. Genes were attributed to a single genomic landscape category in the order listed in figure legend above the plot. (**b**) Enrichments for each category of non-coding SNVs for positive control genes segregated by trait/disease. Enrichment for protein or transcript altering variants was excluded for legibility. (**c**) Enrichment of positive control genes by distance to non-coding SNVs (x-axis) for all traits. Fold enrichment was calculated as the ratio between the proportion of positive control genes targeted to the proportion of all genes targeted. The SNVs with log_10_(BF) > 2 were further overlapped with a master list of DHSs in any cell or tissue type (**Table S5**).

### Generating the *Ei* predictive model

Given the observed enrichment of certain genomic annotations with positive control genes, we next sought to develop a predictive model using the enriched features (described in the **Methods** and summarized in **Fig. 1a**). Briefly, after defining GWAS loci and obtaining a set of fine-mapped SNVs, each SNV was first assessed for a set of annotations, including functional protein-coding or non-coding impact. All annotations are shown in **Table S6**. Then, these annotations were used to map SNVs to genes at each locus (e.g. distance from SNV to each TSS for each gene at the locus; see **Materials and Methods**). Finally, gene-level summary features were developed for all SNVs paired to a given gene to capture both the overall “intensity” of an annotation (e.g. the minimum, mean or maximum) as well as how these intensities varied with distance to gene, measured directly or inversely. This process resulted in a set of primary features used in the prediction model (**Table S7**).

We next took several precautions to ensure the validity and utility of the predictive model. First, since each locus contains features that are shared across all genes at the locus, such as the number of genes at the locus, we incorporated such locus-level features into the model. For instance, if there is only one gene at a GWAS locus, the probability of that gene being causal is higher than if there are a dozen genes at the same GWAS locus. To control for factors associated with locus-wide probability of causality, we included locus-level metrics as features and tested whether the model outperformed these predictors. Second, some diseases and traits studied share GWAS loci and/or positive control genes, such as glucose level and T2D. To avoid cross-contamination of the training and validation sets,[28] the genes were randomly retained for only one trait. Therefore, the same positive control genes were not used for correlated traits, such as glucose level and T2D, thereby preventing over-fitting of the model. Similarly, positive control genes were randomly selected to be included in analysis of only one trait, when multiple traits were involved.

We then trained models using two classifier methods, including logistic regression and XGBoost. The analysis was restricted to loci with at least one positive control gene. The training was conducted using a leave-one-out approach (i.e. leaving out one disease or trait) and the final predictions were then aggregated across all 12 diseases and traits. For comparison with established eQTL-based approaches, we ran eCAVIAR[8] using relevant tissues for each trait from GTEx, and we assessed whether the top posterior probability for a gene across tissues predicted positive control gene status (**Table S5, Materials and Methods**). We recognize that eCAVIAR is meant to identify eQTLs, rather than positive control genes as we have defined them. Nevertheless, this comparison allows for a calibration of performance relative to this known metric.

Aggregating predictions across all 12 traits revealed that XGBoost performed substantially better than logistic regression, achieving an area under the curve (AUC) for the receiver operator curve (AUC-ROC) of 0.79 versus 0.58, and an AUC for the precision recall curve (AUC-PRC) of 0.24 versus 0.09 (**Table 3**). Performance was also higher than eCAVIAR and DEPICT, which achieved an AUC-ROC of 0.71 and 0.63, and AUC-PRC of 0.02 and 0.04, respectively (**Table 3**). These results outperformed simpler approaches, such as selecting the gene nearest the most strongly associated SNV (**Fig. 3**). As a result, we refer to the algorithm for generating features and the prediction model from XGBoost as the Effector index (*Ei*) (**Fig. 3**). These findings suggest that the *Ei* has reasonable prediction accuracy to identify causal genes for common diseases and traits. Disease and trait-specific AUC-ROC and AUC-PRC from the leave one out analysis are shown in **Table S8**.

**Table 3.**
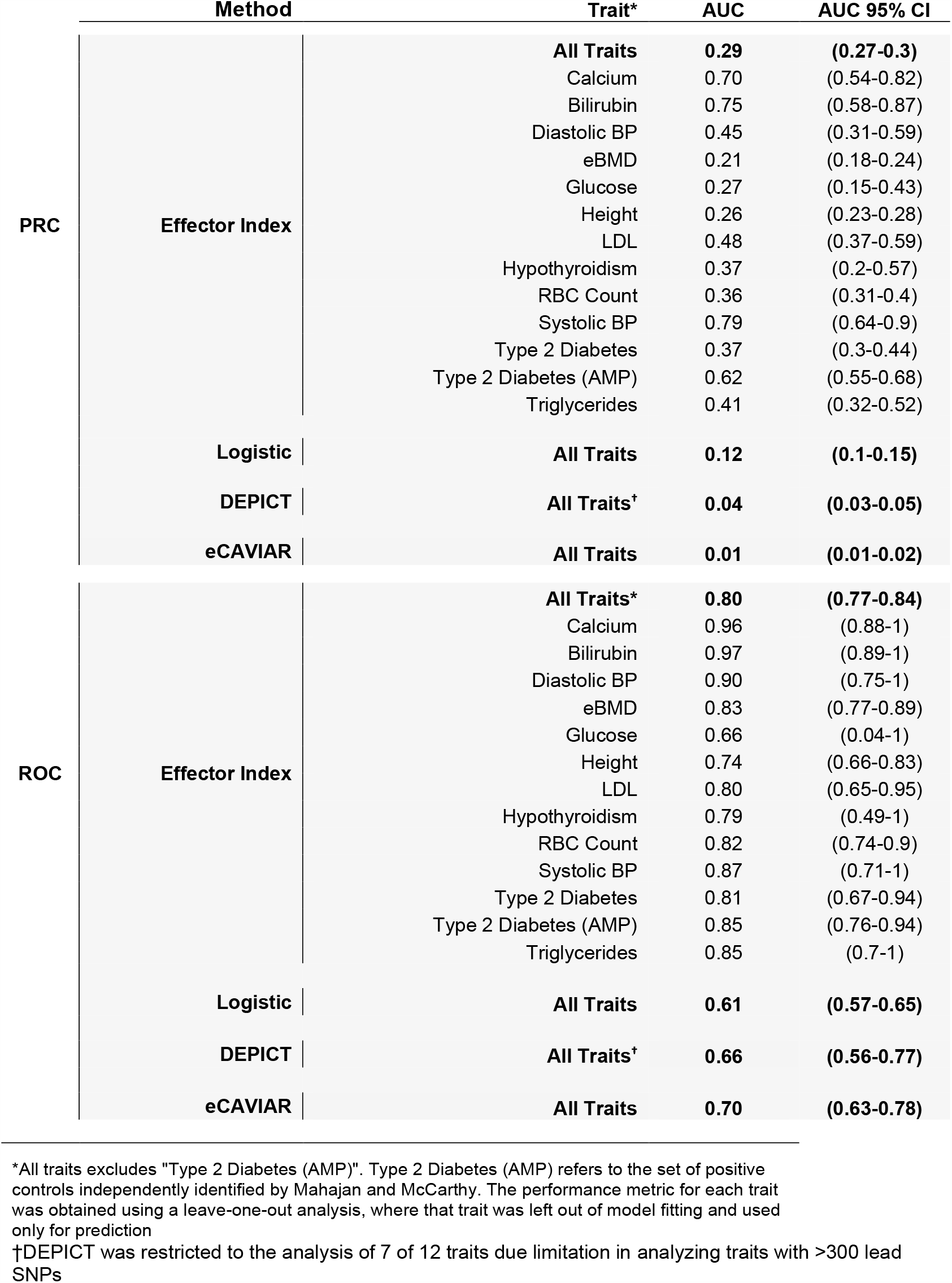
Performance of the Effector index.

**Fig. 3.**
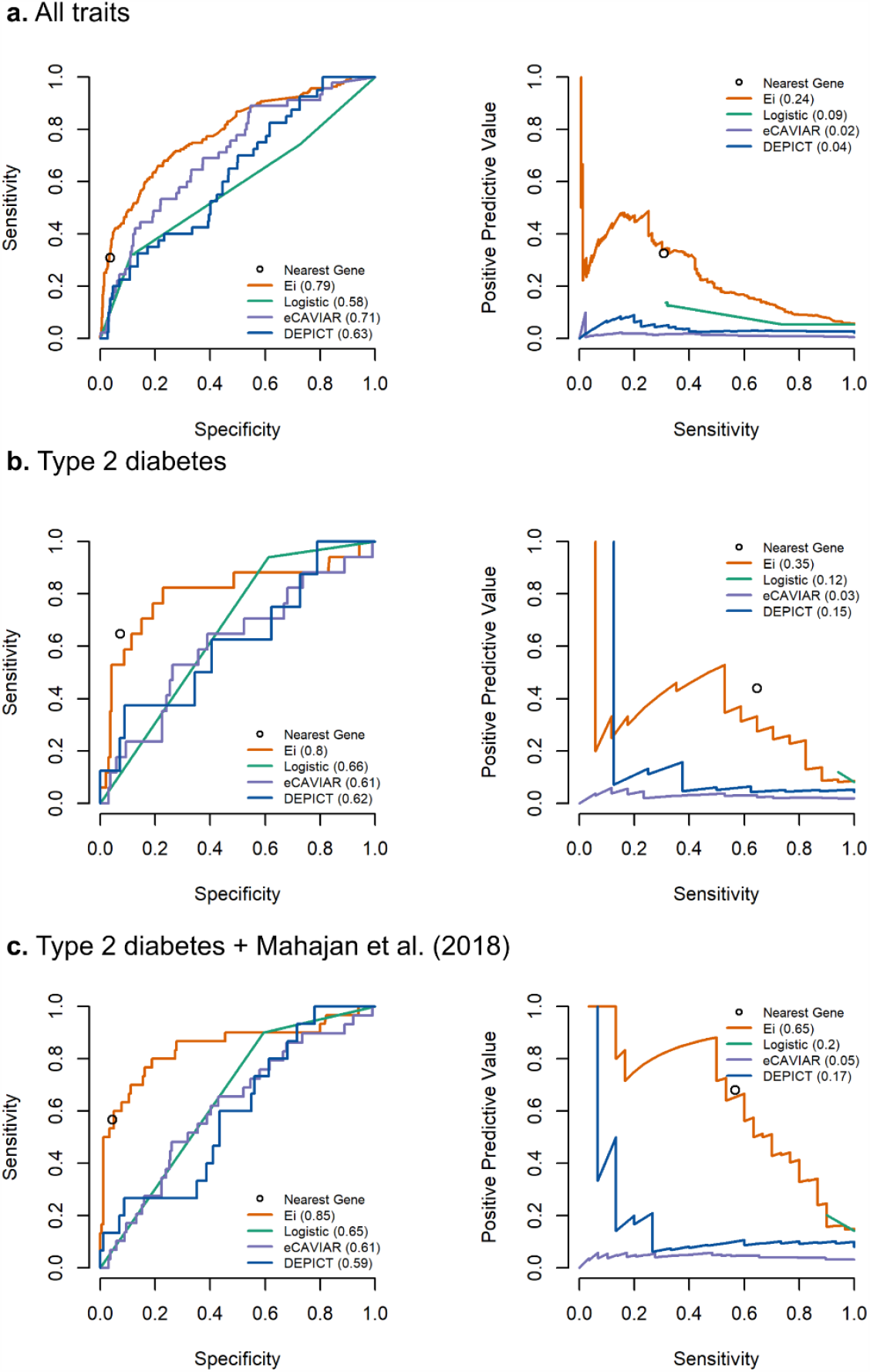
Performance of the Effector index at loci containing positive control genes. The performance of the Effector index compared to logistic regression, eCAVIAR, and DEPICT for predicting positive control genes for: (**a**) all 12 traits, (**b**). type 2 diabetes only, and (**c**) type 2 diabetes with the addition of manually curated causal genes from large-scale exon array and exome sequencing studies.[16] Area under the curves are provided in parentheses and are segregated by trait/disease **Table 3**. Performance using the nearest gene to the lead SNV by P-value is also shown by open circles.

### *Ei* Validation

We next measured the predictive performance of the *Ei* against an augmented set of positive control genes for T2D, which were selected using a complementary approach to select positive control genes. Independently, Mahajan and McCarthy generated a list of 35 positive control genes for T2D[13] (**Table S9**) that included T2D drug targets, genes causing Mendelian forms of T2D, and coding evidence from large-scale exome arrays studies and strong evidence from gene-based burden tests from large-scale whole exome sequencing studies (**Methods Section** and **Table S9**)[29]. The performance of the *Ei* was evaluated in this validation set by first training the *Ei* on all other 11 diseases and traits and testing its performance for T2D using this augmented set of 35 genes as the positive control genes, 24 of which were present within a GWAS locus for T2D as defined in this study.

Using this enlarged and independent list of positive control genes, the AUC-ROC for T2D increased from 0.81 to 0.85 (95% CI: 0.76-0.94) and the AUC-PRC improved substantially from 0.37 to 0.62 (95% CI: 0.55-0.68). These results indicate that in order to have a positive predictive value of ∼80%, the *Ei* would provide a sensitivity of ∼40% to identify positive control genes for T2D (**Fig. 3c**). The *Ei* probabilities of causality between these two sets of positive control genes were similar (**Table S10**). Specifically, 61% of the original positive control genes had an *Ei* probability of >0.8, whereas 63% of the Mahajan and McCarthy positive control genes had an Ei probability of >0.8. This suggests that the *Ei* is able to assign high probabilities to causal genes that are not necessarily known drug targets or causes of Mendelian forms of disease. These findings also demonstrate that as the number of positive control genes identified through large-scale whole-exome sequencing and exome arrays increases, the utility of the *Ei* for mapping GWAS associations to positive control genes is likely to improve. We conclude that applying the *Ei* to data from currently available large-scale GWAS can help to anticipate the results from exome sequencing studies.

### Features influencing the *Ei*’s performance

Given the validation of the *Ei* and its favorable prediction performance, we next asked which features received the largest importance in the model (**Table S7 and Fig. 4)**. Amongst the top 20 features, 4 were influenced by the physical distance of SNVs to genes. The second highest ranked feature was the rank of predicted gene impact from SNPEff (**Table S6** and **Materials and Methods**). Enrichment analysis of these ranked gene impacts, comparing genes containing SNVs with only lowest rank (MODIFIER) to genes with one more SNVs with higher ranks (LOW, MODERATE and HIGH), revealed that higher ranks are more predictive of positive control genes (OR 9.21, p < 2 ×10^−16^). As expected, this demonstrates that coding and transcript altering variation is predictive of causal genes, as these properties are within the higher impact ranks from SNPEff. Conversely, low impact rank is less predictive, as it includes SNVs in introns or within close proximity to the gene and also accounts for the majority of SNVs with SNPEff predictions (**Table S11**). Another predictive feature was fine-mapped SNVs within DHSs in disease-relevant tissue types that are nearest to a gene.

Notably, four locus-level features, such as number of genes at the locus, which results in the same value for all genes at a locus, were also of high importance to prediction (**Fig. 4**). To investigate the ability of the *Ei* to identify positive control genes over and above the performance provided by only locus-level features, we trained models and generated predictions for only the 14 locus-level features (**Table S7**). We found that the *Ei* substantially out-performed locus level features (**Fig. 4b & 4c**), in that models built with only locus-level features provided an AUC-ROC of only 0.73 (95% CI: 0.69-0.77) and an AUC-PRC of only 0.14 (95% CI: 0.08 0.23). We conclude that the *Ei* provides enhanced capabilities to discriminate between genes within a locus.

**Fig. 4.**
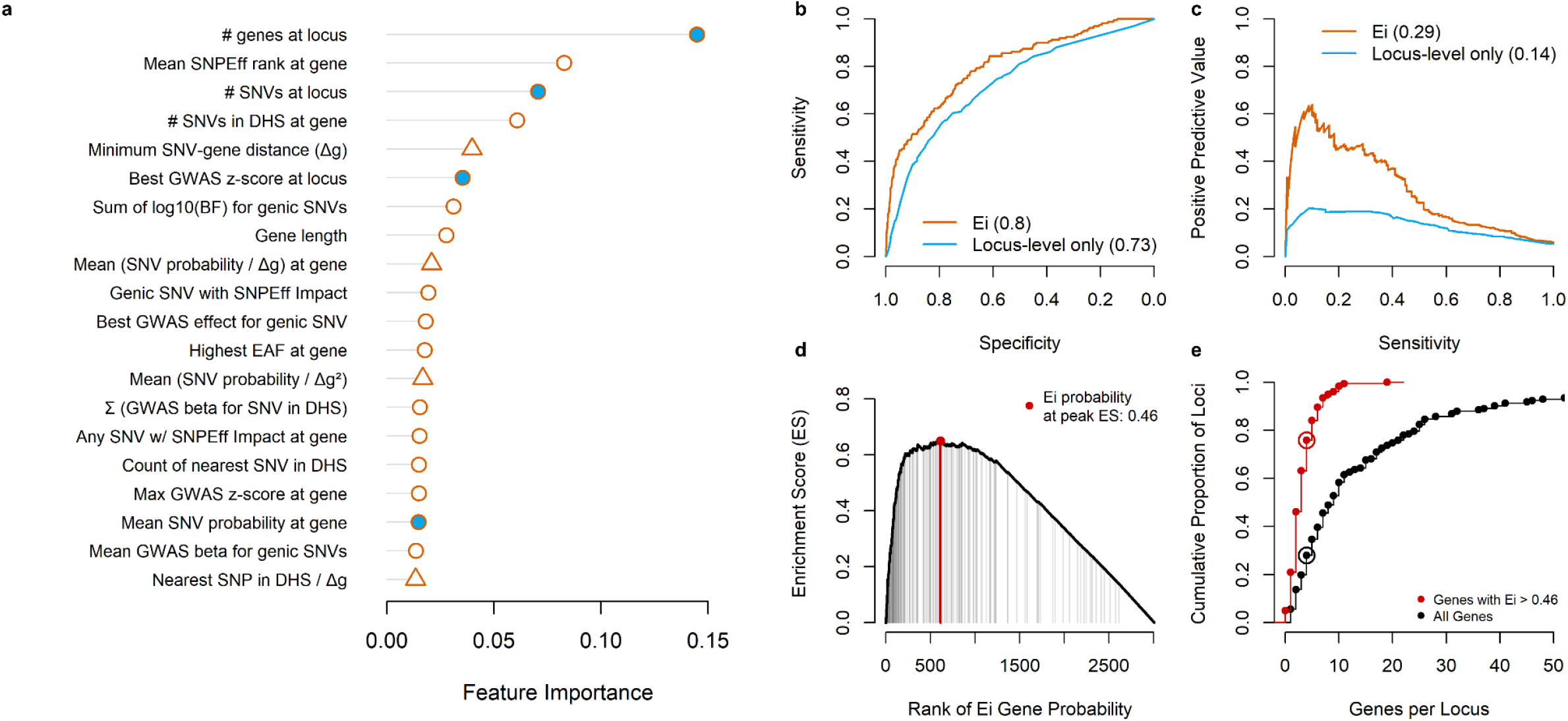
Features selected by the Effector index and comparison of the Effector index to use of only locus-level features. (**a**) Top 20 features selected by the Effector index where each importance value provided is the absolute mean importance of that feature across the 12 traits. Locus-level features shown (in blue) are those do not vary across genes at a locus. Features that incorporate distance to gene are displayed using triangles (Δg denotes SNV-gene distance; ‘genic’ denotes that SNV overlaps gene body). (**b-c**) The ROC (**b**) and PRC (**c**) curves for only locus-level features versus the use of all features in the *Ei* model. Areas under the curve are provided in parentheses. (**d**) Leading edge analysis shows the peak enrichment score for positive control genes occurs an *Ei* probability of 0.46 (red point); a threshold above which we observe 99 of 159 positive control genes (vertical grey lines). (**e**) Using the peak *Ei* threshold of 0.46 considerably reduces the number of genes per locus. For instance, 78% of loci contain 4 or fewer genes with the *Ei* > 0.46 (red open circle), whereas only 28% of loci contain 4 or fewer genes when no threshold is applied (black line open circle

We next sought to test whether the *Ei* could predict a subset of genes at a locus at elevated causal probability, rather than simply assigning the same probability to all genes at a locus. To do so, we first determined the predicted probability cutoff that lead to optimal enrichment of positive control genes and then used this cutoff to select putatively causal genes at each locus. We determined the optimal cutoff to be 0.46, a threshold above which we observe 99 of 159 (62%) positive control genes (**Fig. 4c**).

When applying this cutoff of 0.46 to all loci, we observed that the number of genes considered per locus reduces substantially. Specifically, we found that after applying this cut-off, 78% of loci harbored four or fewer genes. Without such a cutoff, only 28% of loci had four or fewer genes to consider for potential causality. This demonstrates that the *Ei* can enrich for known causal genes and is able to considerably reduce the number of genes to be considered for biological interrogation at a locus.

## Discussion

We have developed the *Ei* which provides helpful predictive accuracy for prioritizing causal genes across a range of traits and diseases. The most useful predictive features were simple metrics such as protein-coding or transcript alterations, fine-mapped SNVs in DHSs and distance from fine-mapped SNVs to genes—features which are often already available in public datasets. We further demonstrated that the *Ei*’s performance increased in a larger and independently curated list of causal genes for T2D, which included genes identified through whole exome sequencing in large cohorts. Importantly, the *Ei* outperforms locus-specific features and simpler algorithms such as nearest gene to the lead GWAS SNV. Last, the *Ei* can substantially reduce the number of genes to be considered for biological validation at GWAS loci. Taken together, the *Ei* provides a publicly accessible algorithm which can be applied to GWAS datasets to enable functional biological dissection and drug development.

A major strength of this study was the use of positive control genes that were derived from Mendelian forms of disease and the targets of clinically useful medicines. While this provides a shorter list of positive control genes than would have resulted from using other methods to establish causality, such as murine or cellular models, it does yield a list of genes whose probability of causality is high. A second major strength is the generalizability of the method across different traits and diseases, influencing different organ systems. Despite the biological differences in the types of conditions studied, the AUC-ROC and AUC-PRC relatively stable.

To improve the generalizability of the *Ei*, we have chosen to focus on a select set of readily available genomic annotations, allowing for its application across many traits and diseases. The annotations we have used can be generated for every GWAS, provided that the imputation reference panel is available to enable accurate fine-mapping and DHS maps from a disease-relevant cell type are available. DHSs have been measured for many tissues and cells via the ENCODE and Roadmap Epigenomics programs, and extension to rarer cell and tissue populations and states is well underway.

We have not used any prior biological information to prioritize genes at loci. While this could be implemented, for example through the use of text-mining tools, we caution that one of the main advantages of the agnostic GWAS approach is the ability to identify genes whose function was not previously known. We note that eCAVIAR, which relies on colocalization of GWAS and eQTL data, did not identify positive control genes as readily as the *Ei*. However, this method is not intended to identify disease causal genes selected through Mendelian inheritance and drug targets, and therefore does not offer a direct comparison. Since our goal is to identify positive control genes, unbiased by prior biological data, we chose to first prioritize genes which can then be complemented by a thorough assessment of the literature.

The relative importance of different predictors in the final *Ei* model is informative. The most important predictor was the simple count of genes at a GWAS locus. This is sensible, since if there is only one gene at a locus, its probability of causality will be higher than if there are 20 genes at the locus. Other informative features were simple metrics such as protein-coding or transcript alterations, distance from fine-mapped SNVs to genes and overlap with DHSs. Previously we have shown that distance to gene is a strong predictor of causal genes in the field of metabolomics,[30] yet there are examples of causal genes at GWAS loci that lie hundreds of kilobases away from the lead SNV[31,32] Further, we have also demonstrated that the nearest gene approach under-performs when compared to the *Ei*, which instead considers proximity in the context of other relevant factors. Given that the majority of positive control genes were associated through non-coding variation, the *Ei* will benefit from improved genomics approaches to infer long-range variant-to-gene links, promising improved prioritization of causal genes for functional dissection and drug development.

This work has important limitations. We have only used large GWASs and emphasize that the accuracy of the *Ei* may not perform well in smaller studies. Further, when implementing fine-mapping, we have been careful to use the same, or very similar reference panels. We would not expect this step of the *Ei* algorithm to perform well if inappropriate reference panels were used. Since fine-mapping strongly enriches for positive control genes, care is warranted when using reference panels different than the original GWAS. While GWAS traits analyzed include various physiologic and metabolic traits, we have yet to fully explore other disease classes, such as neurological traits or diseases. Such future analyses will validate whether the *Ei* model can be further generalized.

In summary, we have developed an implementable algorithm that can identify positive control genes at GWAS loci with reasonable accuracy and can therefore be used to prioritize genes for functional dissection and drug development programs.

## Materials and Methods

A flow diagram that describes the data and process flow of the Effector Index is provided in **Supplementary Fig. 9**.

### Selecting positive control genes

Positive control genes were defined via two approaches: (1) clinician scientists manually inspected the Human Disease Ontology database[17] for relevant ontological terms (**Table S1**), and the associated OMIM linkage information was used to obtain a list of genes associated with these disease (**Table S2**); and (2) clinician scientists identified guideline-approved medications from UpToDate[19] (**Table S3**) and this information was linked to Drugbank[20] to obtain a list of drug targets for drugs with a known mechanism of action (**Table S4**). The above procedure was performed for all traits, except for eBMD, for which we extracted the positive control gene list from Supplementary Table 12 of Morris et al. 2018.[12] To assess the performance of the prediction models, we also used data from the National Institutes of Health T2D Accelerated Medicines Program, a collaboration between industry and academia to identify causal genes for disease.[16] We included the original T2D genes described identified as monogenic causes or drug targets for T2D as above and included additional genes labeled to be “causal”. Such additional genes met any of the following criteria: 1) Exome array evidence with a cumulative posterior probability of association (PPA) of ≥80%;[33] 2) Burden test evidence from WES, using the best gene-level p-value from the extreme p-value aggregation test, or weighted aggregation test performed in an exome sequencing analysis of over 49,000 individuals[34] and had a P-value 5×10^−6^; 3) Strongly associated coding variants reported in the literature.[35] We removed all genes that were labeled as “causal” due to evidence only from GWASs, as including such genes would bias our algorithm’s performance away from the null.

### Obtaining GWAS summary statistics and defining associated loci

GWAS summary statistics were obtained from a combination of publicly available resources and though the GWAS of UK Biobank traits in this study (**Table 2**). These diseases and traits were T2D, LDL levels, red blood cell count, diastolic blood pressure, systolic blood pressure, triglyceride levels, estimated bone mineral density, glucose levels, calcium levels, direct bilirubin levels, height and hypothyroidism. These traits were selected because they represent a broad spectrum of allelic architectures, ranging from oligogenic to highly polygenic as evidenced by the variable number of loci identified using the method described below (**Table 2**).

For the GWAS of traits generated in this study we used the White-British subset of individuals in UK Biobank (N=440,346); an analysis that was performed in a previous study that consisted of the projection of UK Biobank individuals to 1000 Genomes, followed by a cluster analysis to identify a subset of individuals of relatively homogenous ancestry.[12]

For the GWAS of bilirubin we natural log transformed the direct bilirubin measurement (UK Biobank data-field 30660) and retained measurements within 3 standard deviations of the mean. For the GWAS of calcium (UK Biobank data-field 30680) we retained unadjusted measurements within 3 standard deviations of the mean. For the GWAS of glucose (UK Biobank data-field 30740) we retained unadjusted measurements within 2.5 standard deviations of the mean. For the GWAS of low-density lipoprotein (UK Biobank data-field 30780) and triglycerides (UK Biobank data-field 30870) we adjusted the measurements for participants taking medications as follows. Participants that self-report taking relevant medications were obtained from UK Biobank data-field 20003 as well as data-fields 6153 and 6177. In reference to previous randomized control trials[36,37] and meta-analyses[38– 41] on the percentage effect of each commonly used medication, we have generated adjustment factors listed in **Table S12**. To estimate the LDL or TG level without medication’s influence, an individual’s measured level was divided by (1 - adjustment factor). Those who did not specify the type of cholesterol-lowering medication were assumed to be on statin, and their adjustment factor was defined as the weighted mean percentage effect of the five statins by their prevalence in UK Biobank population. Further details of the adjustments are provided in **Table S12**.

The GWAS for the 5 traits was performed using fastGWA[42] on SNVs with minor allele frequency greater than 0.0001 and information score (INFO) greater than 0.8. Age, sex, assessment center, genotype array, and first 10 principal components of ancestry (PCs) of ancestry (derived from the principal component analysis in the white British cohort in UK Biobank) were used as additional covariates.

GWAS summary statistics across the 12 traits were harmonized by retaining only SNVs with minor allele frequency below 0.005 that were also present in UK Biobank (matched by SNV alleles and genomic position to GRCh37). The harmonized SNV count for each trait is listed in **Table 2**.

### Defining GWAS loci for Bayesian fine-mapping

GWAS loci were defined using a two-step procedure of LD clumping followed by merging of adjacent signals. First, we determined a set of lead SNVs by LD clumping genome-wide significant SNVs using PLINK 1.9 (p < 5 × 10^−8^, *r*^*2*^ < 0.01, distance 250 kilobases) to a reference panel of 50,000 randomly selected white British individuals (N=409,703) as determined by Bycroft et al. (2018)[43]. Second, lead SNVs that were within 50 Kbp of each other were merged using bedtools ‘merge’. The resulting genomic regions consisting of one or more lead SNVs were padded with an additional 250 Kbp, resulting in a set of loci that are at least 500 Kbp in size, but some loci may be larger due to multiple adjacent lead SNVs. Loci that overlap the major-histocompatibility complex locus were excluded (chr6:28477797-33448354 on GRCh37).

### Finding causal SNVs using Bayesian fine-mapping

We used Bayesian fine-mapping as implemented by FINEMAP version 1.3.1 to find a set of putatively causal SNVs at each locus. This program uses a shotgun stochastic search algorithm to efficiently search for possible causal configurations of up to *k* SNVs at a locus. We used this program to find causal SNVs at a locus (up to a maximum of *k*=20) by providing as input the GWAS summary statistics at the locus, as well as the same reference panel of 50,000 individuals from UK Biobank that was used for defining the GWAS loci in the previous section. As recommended by Benner et al. (2017)[44], the population of this reference panel matches what we used for the UK Biobank GWAS or is similar to the publicly available GWAS included in this study. We are unaware of any other method that efficiently searches for causal configurations containing up to 20 causal SNVs and across as many traits as we have explored in this study, allowing us to fine map large loci with potentially complex signals.

To do this, we created a simple shell script that extracted individual level genotype data from UK Biobank using [bgenix](https://bitbucket.org/gavinband/bgen/wiki/bgenix) and from this calculate the genotype correlation matrix using LDStore.[44] GWAS summary statistics were formatted for input into FINEMAP and the prior standard deviation of effect size was determined for case-control studies as defined by Benner et al. (2016) (**Table S13**).[21] Loci that report a sum of posterior probabilities of < 0.95 for the causal configurations 19 SNVs or less were deemed to have failed convergence and were discarded.

The GWAS loci defined for Bayesian fine-mapping may overlap due to lead SNVs being beyond the maximum distance allowed for locus merging (50 Kbp), but the within the padded distance added to each locus of 250 Kbp. As a result, this generated multiple FINEMAP summary statistics for a SNV in a region overlapped by multiple GWAS loci. For these SNVs, we conservatively assigned the FINEMAP summary statistics that report the lowest log_10_(BF). As a consequence of this harmonization, we were then able to merge overlapping loci used for Bayesian fine-mapping into a single larger locus. SNVs achieving a log_10_(BF) > 2 were retained for further analyses as this threshold is generally considered to be strong evidence for causality,[45,46] and we have previously shown that SNVs at or above this threshold are enriched for missense SNVs and of SNVs at accessible chromatin sites.[12]

The number of loci, GWAS lead SNVs, and SNVs with log_10_(BF) > 2 that were retained per trait for subsequent analyses are listed in **Table 2**.

### Source or generation of DNase-seq data

Saos-2 and U2OS cells were maintained in adherent cultures in McCoy’s 5A medium supplemented with 1x Penicillin/Streptomycin and Fetal Bovine Serum (FBS) (15% and 10%, respectively). Saos-2 and U2OS cells were sub-cultured at a ratio of 1:3 and 1:6, respectively, once they reached 80% confluency. DNase digestion was performed as described previously[47] adapted to 200 uL thermocycler tubes. Briefly, nuclei were extracted from cells and incubated with limiting concentrations of the DNA endonuclease DNase I (Sigma) supplemented with Ca^2+^ and Mg^2+^ for 3 min at 37 °C. Digestion was stopped by the addition of EDTA, and the samples were treated with proteinase K and RNase A. Short double hit fragments from DNase digestion using magnetic bead polyethylene glycol (PEG) fractionation. Illumina libraries were generated and sequenced on an Illumina NextSeq 500.

DNase-seq datasets from the ENCODE and Roadmap projects were downloaded from http://www.encodeproject.org.[48,49] ATAC-seq data of pancreatic islets was downloaded from SRA SRR8729334.[27] See also **Table S5**.

All DNase-seq and ATAC-seq data were processed using a uniform mapping and peak calling pipeline (https://github.com/mauranolab/mapping/tree/master/dnase). Illumina sequencing adapters were trimmed with Trimmomatic.[50] Reads were aligned to the human reference genome (GRCh38/hg38) using BWA.[51] Hotspots were called using hotspot2 (https://github.com/Altius/hotspot2) with a cutoff of 5% false discovery rate. Hotspots were converted to hg19 reference coordinates using UCSC liftOver. Saos-2 and U2OS DNase-seq data are available from GEO at accession GSE142160.

### Tissue-selective Expression Positive Control Gene Sets

Tissue-selective expression was established by differential expression analysis in a selection of 32 tissues from the GTEx project v7 obtained from the GTEx Portal on 09/27/2017.[49] The top five RNA-seq samples in terms of data quality were selected per tissue, and all pairwise differential expression analyses were performed using DESeq2.[52] A gene was considered differentially expressed between two tissues if it passed cutoffs for both log_2_(fold change) > 3 and Bonferroni-adjusted P-value < 0.01. Bonferroni-adjusted P-values were calculated to account for all pairwise comparisons. Gene sets for each tissue were defined as genes differentially expressed in at least 26 (50%) comparisons. See also **Table S14**.

Processed transcript quantification[53] of RNA-seq data was downloaded for purified T-cells, B-cells and monocytes (CD4 or CD8, CD14, and CD20) from ENCODE[48,49,54]; http://encodeproject.org, accessions ENCFF269QBU, ENCFF880QDD, ENCFF557VGS, ENCFF495CNV, ENCFF081MXC, ENCFF669GZO, ENCFF049GRB, and ENCFF422SXS). Gene sets were generated as above, except that a differential expression cutoff of log_2_(fold change) > 2 was used. Gene sets for each tissue were defined as genes differentially expressed in at least 2 (50%) comparisons.

For T2D, additional gene sets for were obtained from RNA-seq of pancreatic islets[55] or scRNA-seq of pancreatic endocrine cells.[56]

All gene IDs were mapped to Gencode v24. Gene lists of similar tissues were merged together (**Table S5**).

### Transcript Annotation

For Fig. 2, SNVs affecting missense coding sequence or transcript structure were identified using the ENSEMBL Variant Effect Predictor v92[57] and Gencode v24, using the following tags: splice_donor_variant, splice_acceptor_variant, stop_gained, stop_lost, start_lost, missense_variant, splice_region_variant, incomplete_terminal_codon_variant, stop_retained_variant, coding_sequence_variant, mature_miRNA_variant, 5_prime_UTR_variant, 3_prime_UTR_variant, NMD_transcript_variant.

### Hi-C and eQTL data

Published pcHi-C data obtained for a subset of traits (**Table S14**).[58–60] Interactions were removed if either contact region was >30 Kbp in length, or if the distance between the midpoints of interacting contact regions was > 1 Mbp. eQTL data v7 was downloaded from the GTEx Portal on 09/27/2017.[49] For each trait, a set of relevant tissue types was defined and all eQTL-eGene pairs passing a 5% FDR cutoff were used (**Table S14**).

### Variant-to-Gene Enrichment

We assessed a variety of criteria to link associated variants to genes through enrichment of significant SNV sets for positive control genes. The background was defined as protein coding and lincRNA genes in the Gencode v24 basic annotation. Enrichments were defined as the proportion of positive control genes within the targeted set divided by the background proportion of positive control genes. All SNVs within coding regions were removed before gene targeting.

### SNV-gene annotation

Putatively causal SNVs from the Bayesian fine-mapping analysis were annotated for potential functional effect using the following methods:

1. Predicted functional impact was extracted using SNPEff version 4.3T for build GRCh37.75 with default parameters and databases (snpEffectPredictor, nextProt, pwms, and protein interactions). The predicted impact annotation was then numerically ranked as follows: 1:’HIGH’, 2:’MODERATE’, 3:’LOW’, 4:’MODIFIER’, 5:’NONE’ (**Table S6**). Note that the MODIFIER impact prediction includes genomic regions outside the gene body, such as regions 5 Kbp upstream and downstream of the gene (see http://snpeff.sourceforge.net/SnpEff_manual.html).
2. We extracted the predicted functional effect of each SNV from version 150 of dbSNP obtained from the UCSC Genome Browser (build GRCh37).
3. We identified any overlap of putatively causal SNVs with one or more DHSs from a set of trait-matched cell/tissue types and to the entire set of 160 cell/tissue types collected from ENCODE and other sources (**Table S5**). See also *“Source or generation of DHS data”*.

Putatively causal SNVs and their annotations were assigned to protein coding genes using GENCODE v29 to identify genes overlapping each locus by ≥ 50% of their length as follows:

1. For each gene at a locus, assign all overlapping putatively causal SNVs.
2. For each gene at a locus, assign the putatively causal SNV nearest to its transcription start site.
3. Assign a putatively causal SNV to gene if the affected gene reported by SNPEff is a gene at the locus (i.e. protein or transcript-altering SNVs).
4. For each gene at a locus, assign the closest putatively causal SNV overlapping a DHS.

Each SNV-gene assignment was additionally annotated with distance from the SNV to the gene TSS, to the gene body, and to the transcription end site (**Table S6)**.

SNV-gene annotation was summarized per gene using average, minima, and maxima to create 140 candidate gene-level features (**Table S7**). We also generated a set of locus-level features which do not vary across all genes at each locus (**Table S7**). Uninformative features were removed, such as those with a standard deviation of 0, and correlated features were consolidated.

### Building and testing predictive models

Model training was performed at loci containing at least one positive control gene. For each trait or disease (*k*), we coded gene (*j*) as a positive control (*Y*_*jk*_) as 1 if the gene was labeled as a positive control gene, or 0 if not. To avoid overfitting, we pruned the matrix so that each gene contributed at most once to model building using the following approach:

1. If a gene is a positive control gene for multiple traits, then one of these traits was randomly chosen and retained for analysis (*Y*_*jk*_ = 1) while the other entries were dropped.
2. If a gene is a positive control gene for some traits, but a negative gene for other traits, then that gene was retained only for the true positive trait (*Y*_*jk*_ = 1). One trait was randomly selected from among the positives, if there was more than one.
3. If a gene is negative (*Y*_*jk*_ = 0) for multiple traits, then one trait was randomly chosen and other traits were dropped.

Training and performance assessment was performed by combining genes across all but one of the traits, and then testing on the trait left out. As each gene can appear a maximum of once, no genes can overlap between the training and testing sets.

We predicted the true causal status of a gene (***Y ∼ X***) by using the gradient boosted trees algorithm, as implemented in the XGBoost package in R (https://cran.r-project.org/web/packages/xgboost/). The input variables (***X***) were standardized prior to statistical modeling and the response variable is binary (***Y = 1*** or ***0***). Each observation corresponding to a negative gene was down-weighted according to the ratio of the positive genes to negative genes in the training dataset for trait *k*. There were few positive control genes relative to negative genes, and therefore this weighting ensures that their features contribute substantially to the model. The hyperparameters in XGBoost, such as tree depth, lambda and gamma, were tuned to optimize the cross-validated performance on training data. Performance was measured using the area under both receiver operator curves (AUC-ROC) and precision-recall curves (AUC-PRC) in the test datasets.

### eCAVIAR

We used eCAVIAR version 2.2 to prioritize genes at each locus by determining the SNVs that are responsible for both the GWAS and eQTL signals. To ensure a comparable prediction performance to Ei we prepared input data as follows: For each trait we selected eQTL GWAS summary statistics for tissues from GTEx v7 as defined previously (**Table S5**). We then obtained trait GWAS summary statistics and genes from loci as defined previously for this study (see “**Defining GWAS loci for Bayesian fine-mapping**”). SNV summary statistics were then harmonized between the trait and eQTL GWAS to include only SNVs matching by NCBI dbSNP identifier and alleles, as well as retaining only SNVs with MAF > 0.01 from GTEx v7. Linkage disequilibrium was computed using the same reference panel described previously for this study (see “**Defining GWAS loci for Bayesian fine-mapping**”). We then used eCAVIAR with one maximum causal SNV per locus (-c 1) to obtain a listing of SNV-gene pairings and the colocalization posterior probability (CLPP) of the SNV being responsible for the eQTL tissue and trait GWAS signals. We then collated this information across trait-tissue pairings and report the maximal CLPP for a each gene across all tissues for that trait.

Specific ethics approval was not required for this study.

## Acknowledgements

The funding agencies had no role in the design, implementation or interpretation of this study. The views expressed in this article are those of the author(s) and not necessarily those of funders. EF is an employee of Pfizer. MIM is funded by the NHS, the NIHR, and the Department of Health. MIM has served on advisory panels for Pfizer, NovoNordisk and Zoe Global, has received honoraria from Merck, Pfizer, Novo Nordisk and Eli Lilly, and research funding from Abbvie, Astra Zeneca, Boehringer Ingelheim, Eli Lilly, Janssen, Merck, NovoNordisk, Pfizer, Roche, Sanofi Aventis, Servier, and Takeda. As of June 2019, MIM is an employee of Genentech, and a holder of Roche stock. MIM has received funding from the NIH: U01-DK105535 and the Wellcome Trust: Wellcome: 090532, 098381, 106130, 203141, 212259. The Greenwood lab acknowledges support from Compute Canada (RAPI: nzt-671-aa). MTM is partially funded by National Institutes of Health grant R35GM119703. The Richards research group is supported by the Canadian Institutes of Health Research (CIHR), the Lady Davis Institute of the Jewish General Hospital, the Canadian Foundation for Innovation, the NIH Foundation, Cancer Research UK and the Fonds de Recherche Québec Santé (FRQS). JBR is supported by a FRQS Clinical Research Scholarship. JBR has served as an advisor to GlaxoSmithKline and Deerfield Capital. TwinsUK is funded by the Welcome Trust, Medical Research Council, European Union, the National Institute for Health Research (NIHR)-funded BioResource, Clinical Research Facility and Biomedical Research Centre based at Guy’s and St Thomas’ NHS Foundation Trust in partnership with King’s College London. This research has been conducted using the UK Biobank Resource using project number 27449.

## Data availability statement

All accession codes and URLs for publicly available data are provided in the **Materials and Methods**. Newly generated DNase-seq data are available from the GEO repository under accession GSE142160. The data that support the findings of this study are available from GitHub at https://github.com/richardslab/Ei. This includes raw data underlying the figures. Results can be visualized at http://hugeamp.org/effectorgenes.html.

## Code availability statement

The code that supports the findings of this study are available from GitHub at https://github.com/richardslab/Ei and https://github.com/mauranolab/UKBB_FINEMAP_targetgene.

