## Supplemental Figures for "An Effector Index to Predict Causal Genes at GWAS Loci"

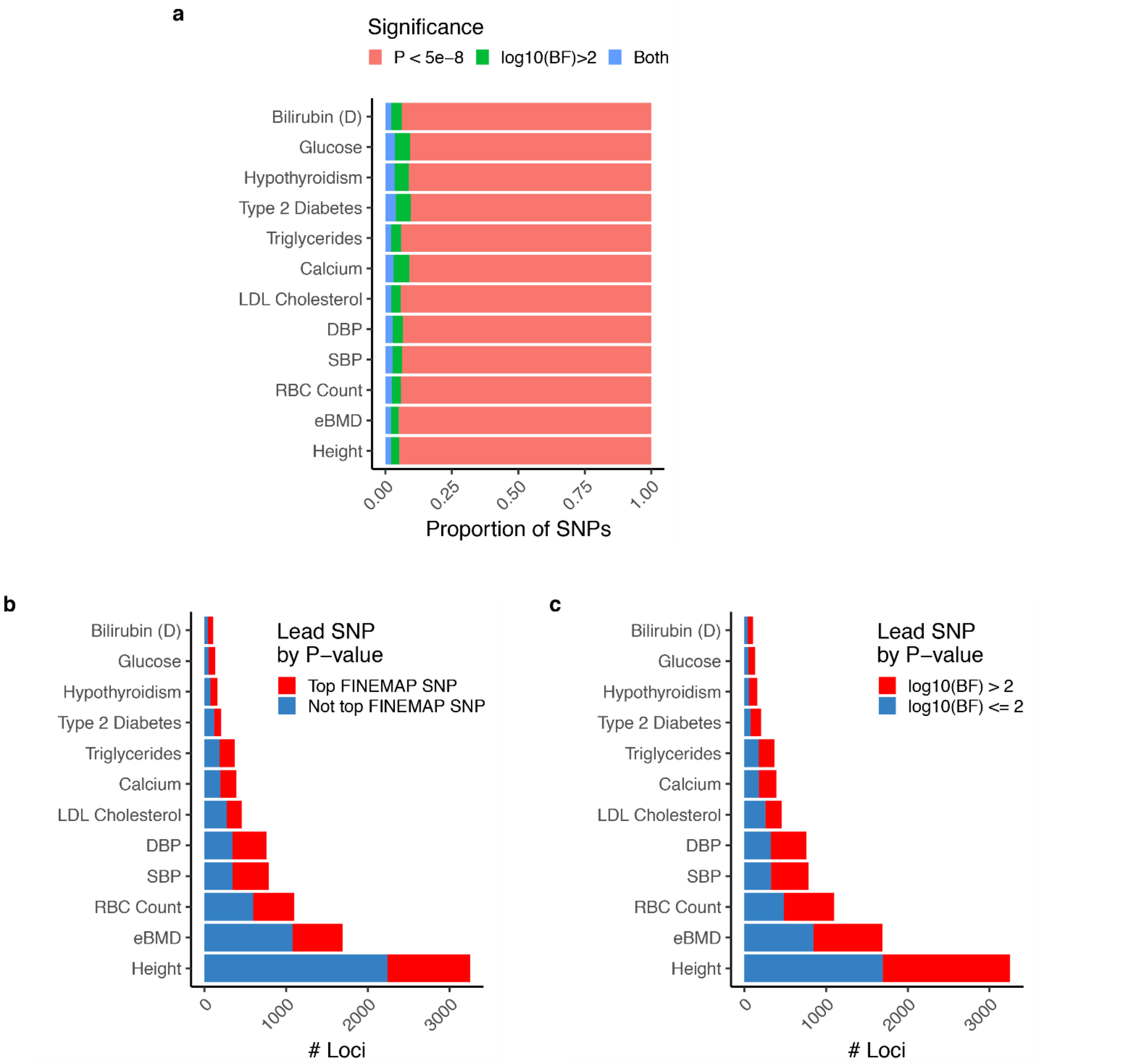


**Supplementary Fig. 1. Summary of statistical fine mapping results.**
(**a**) Composition and overlap of genome-wide significantly associated sets and FINEMAP credible set. (**b-c**) Frequency of loci where lead GWAS SNV by P-value is also (**b**) lead SNV by $\log_{10}$(BF) or (**c**) has $\log_{10}$(BF) > 2.

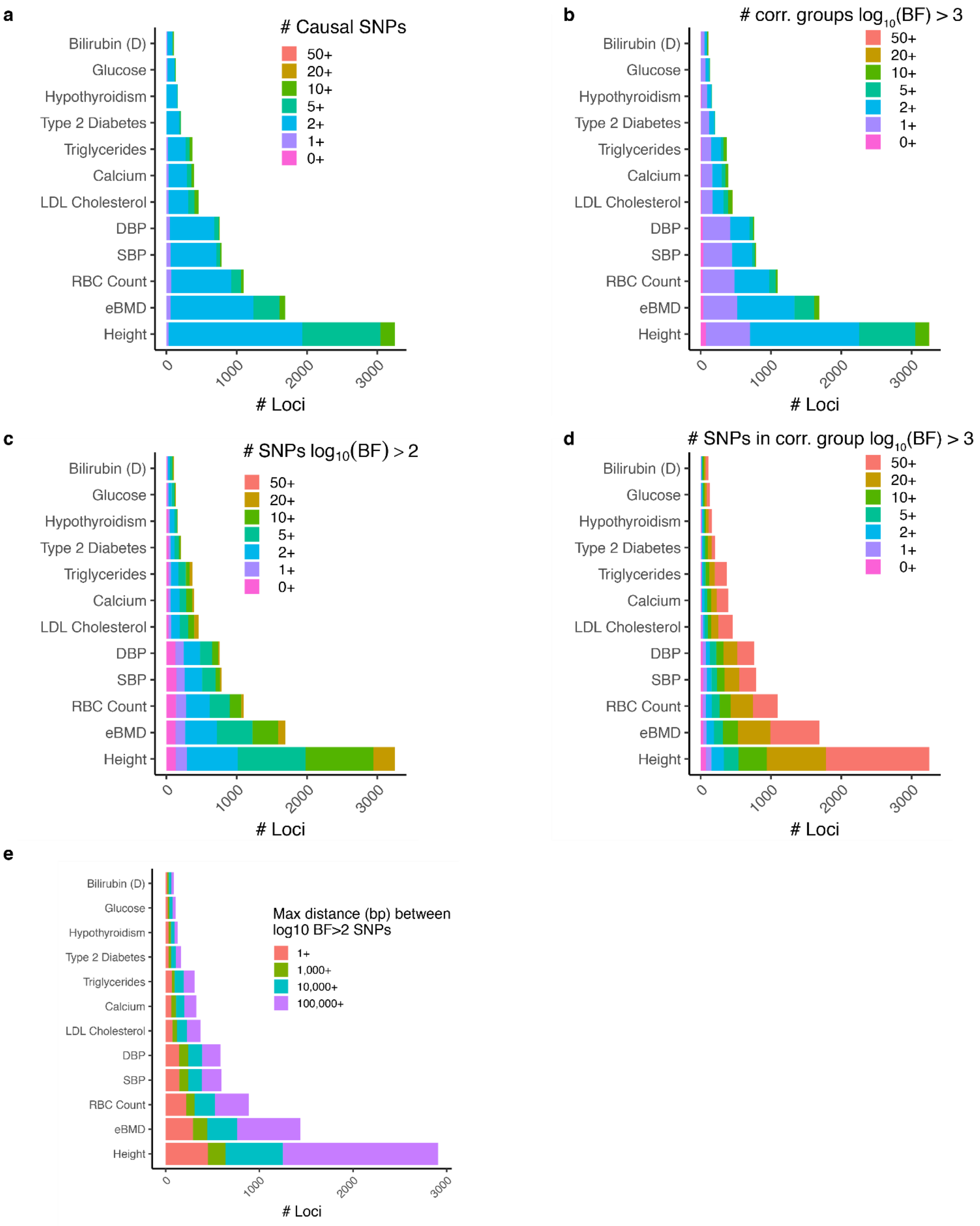


**Supplementary Fig. 2. Assessment of fine-mapping GWAS loci.**

Overview of allelic heterogeneity at fine-mapped loci (**a-d**). Accounting for the number of causal signals per trait, as measured by (**a**) FINEMAP (k_max); (**b**) number of statistically significant groups of independent SNVs $\log_{10}(BF) > 3$; (**c**) the number of individual SNVs $\log_{10}(BF) > 2$; and, (**d**) the number of SNVs in statistically significant groups of independent SNVs $\log_{10}(BF) > 3$. (**e**) extent of genomic region covered by $\log_{10}(BF) > 3$ at each locus.

**a**

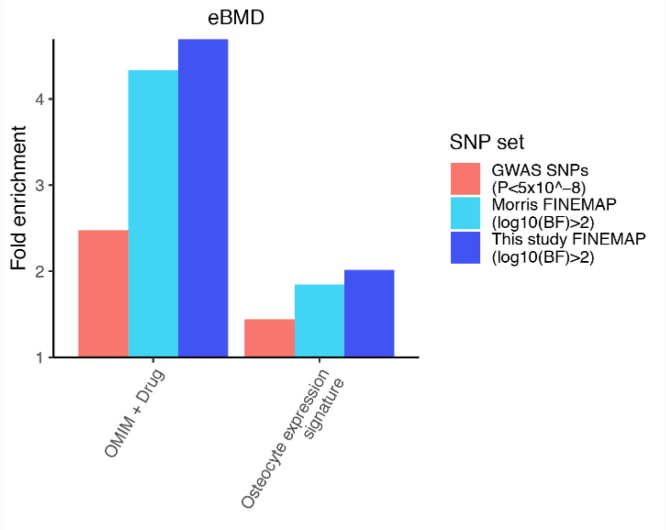


**b**

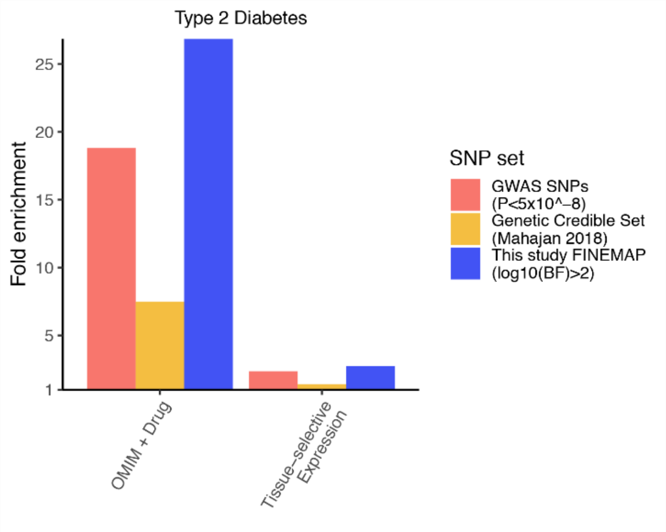


### Supplementary Fig. 3. Comparison of fine-mapping credible sets to past studies.

(**a-b**) Enrichment of positive control gene sets. Shown is enrichment of SNV sets for positive control genes within 25 Kbp. (**a**) Comparison to eBMD analysis from Morris et al. 2019[1] in terms of enrichment for OMIM + Drug genes and osteocyte expression genes. The Morris et al. GWAS was used as input to the FINEMAP pipeline in the present study. (**b**) Comparison to published Type 2 Diabetes fine-mapping study[2] in terms of enrichment for OMIM + Drug genes and Expression positive control sets.

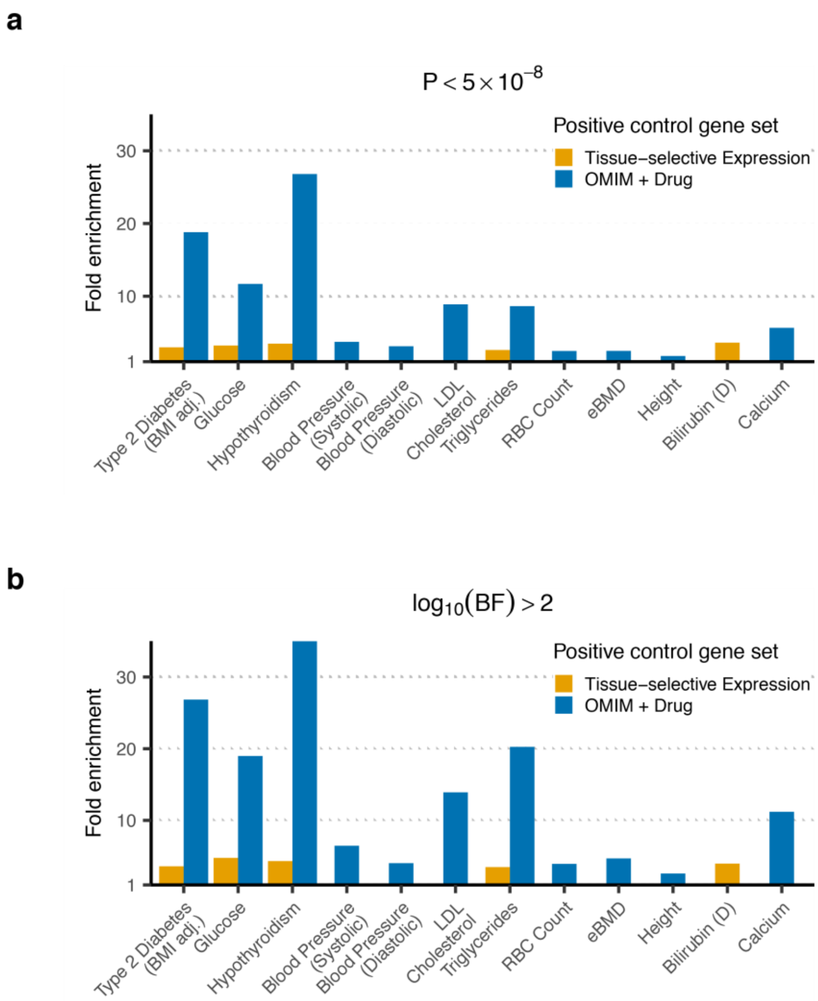


**Supplementary Fig. 4 Fine mapping yields increased enrichment for positive control genes.** Comparison of fold enrichment for positive control genes within ±25 kbp of (**a**), significant ($P < 5 \times 10^{-8}$) or (**b**), $\log_{10}(BF) > 2$ SNVs. Fold enrichment was calculated as the proportion of positive control genes targeted to the proportion of all genes targeted.

a

b

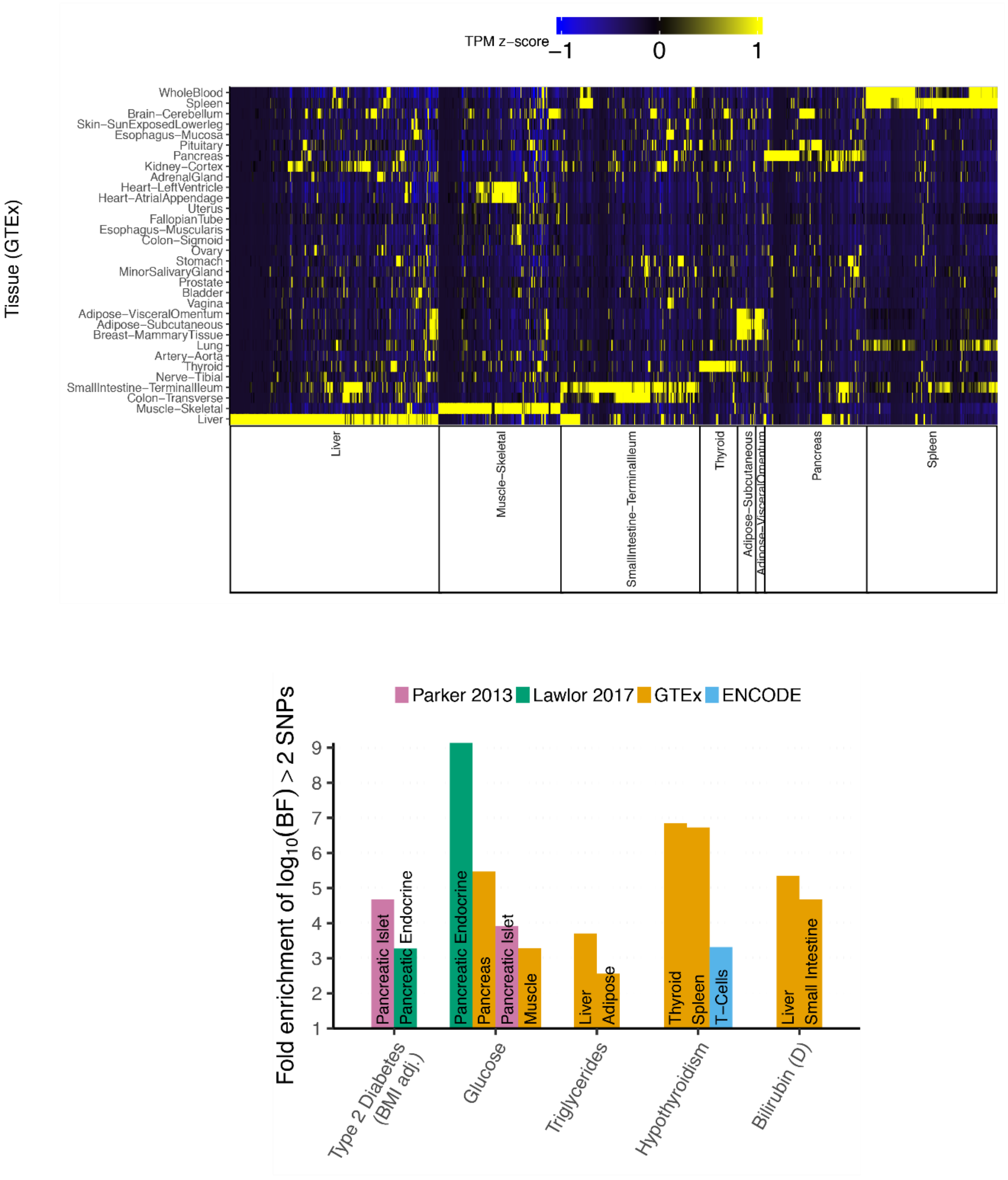


**Supplementary Fig. 5. Tissue-selective expression positive control gene sets.**
(**a**) Expression of genes by RNA-seq in relevant tissue-selective sets (x-axis) for 32 tissues from GTEx (y-axis). Color indicates TPM level normalized by z-score across tissues.
(**b**) Enrichment for tissue-selective gene sets within 25 Kbp of $\log_{10}(BF) > 2$ SNVs for 5 traits.

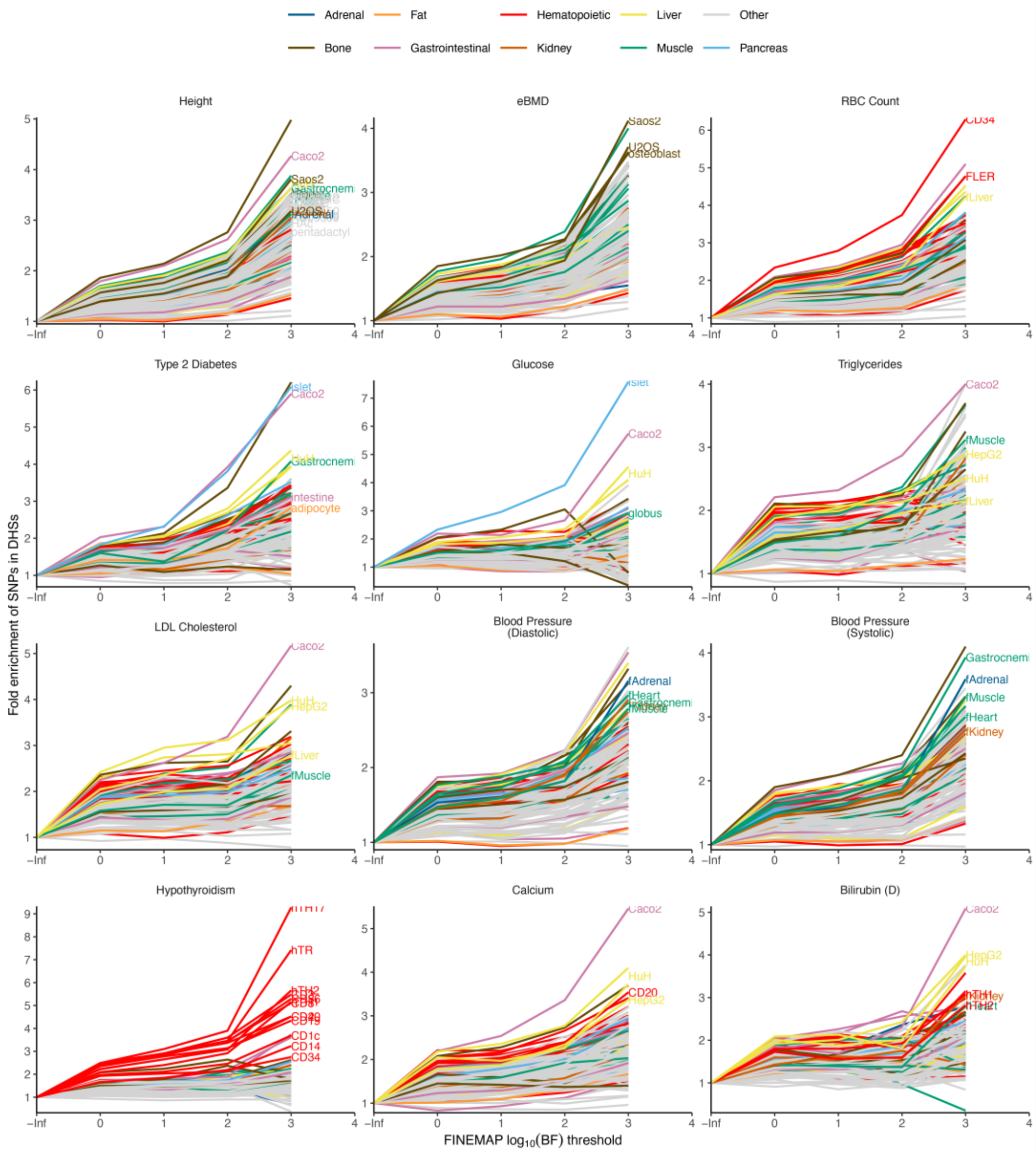


**Supplementary Fig. 6 Enrichment of SNV sets meeting increasing $log_{10}$(BF) cutoffs for DHSs.** Labeled enrichments correspond to cell and tissue types selected for that trait. Enrichments were calculated as the ratio between the proportion of SNVs above threshold within DHSs and the proportion of all SNVs within DHSs.

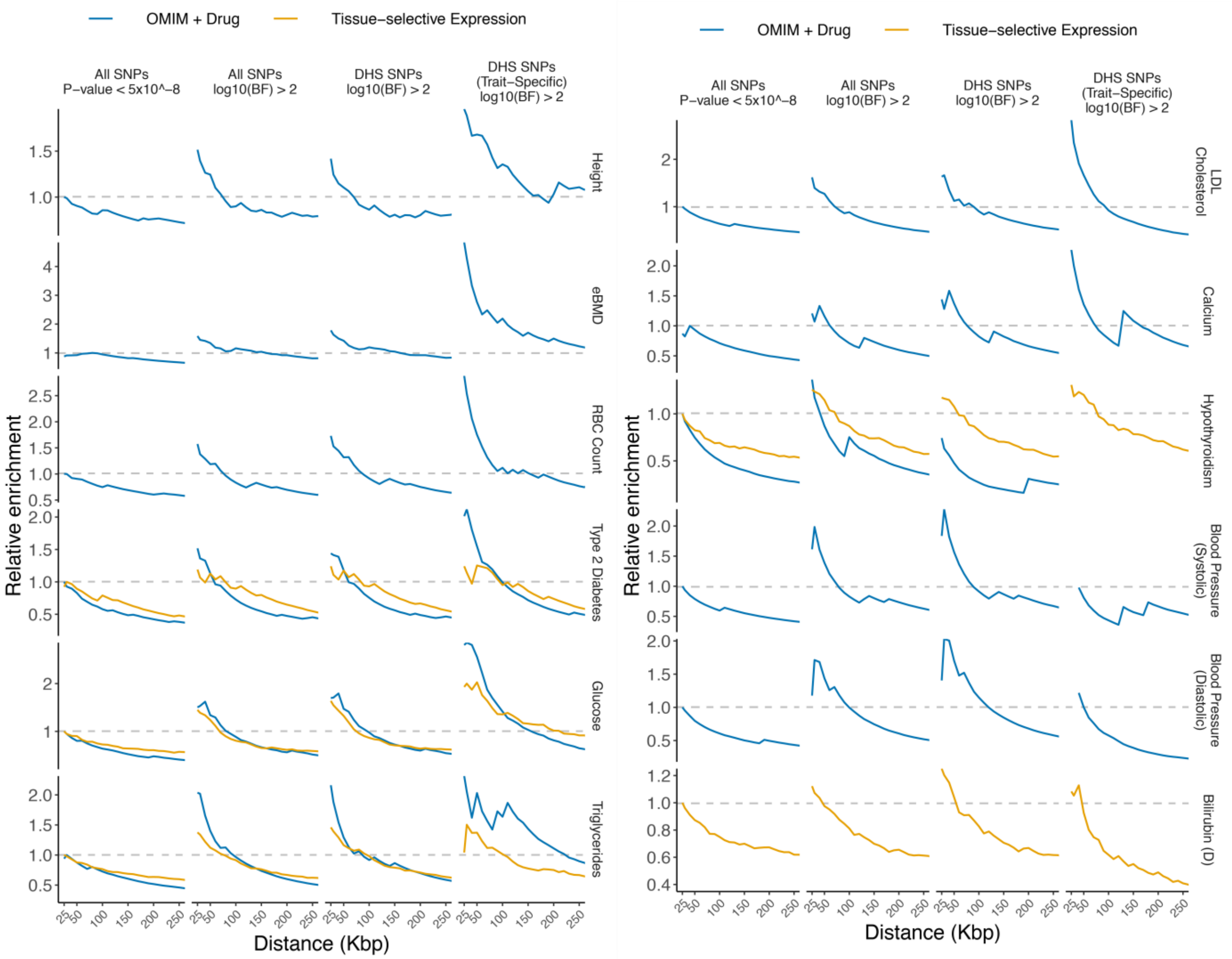


**Supplementary Fig. 7. Gene targeting enrichment (normalized).**
Gene targeting enrichment over distance (x-axis) for all traits. SNV sets, DHS sets, and enrichments are as in **Fig. 2c**, but normalized to the peak enrichment in “All SNVs, P-value < $5 \times 10^{-8}$.” The OMIM + Drug set for Bilirubin (D) has only 2 genes and is not shown.

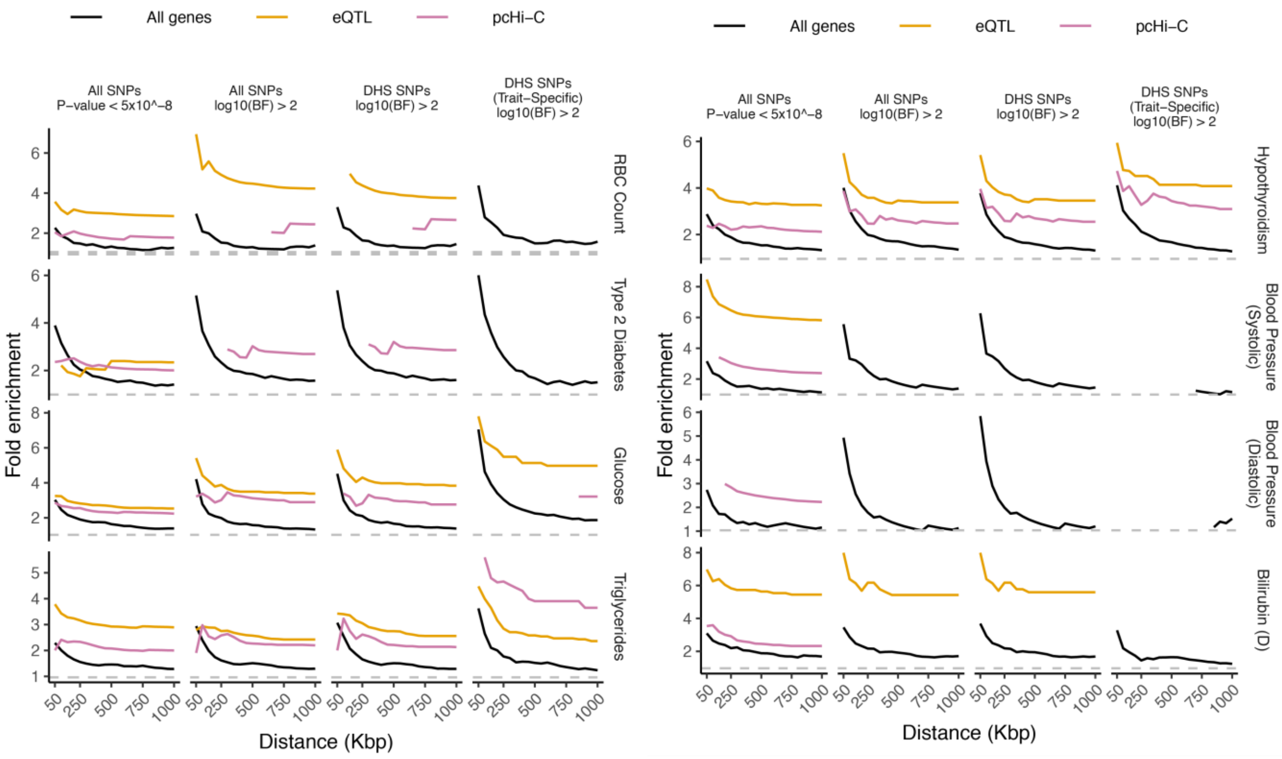


**Supplementary Fig. 8. Long-range gene targeting using pcHi-C and eQTL data.**
Enrichments by distance shown using GTEx eQTLs and pcHi-C (**Table S5**). SNV sets and enrichments are as in **Fig. 2c**. OMIM + Drug and Expression gene sets were combined to serve as positive controls. Enrichments are shown for points where >3 positive control genes are represented.

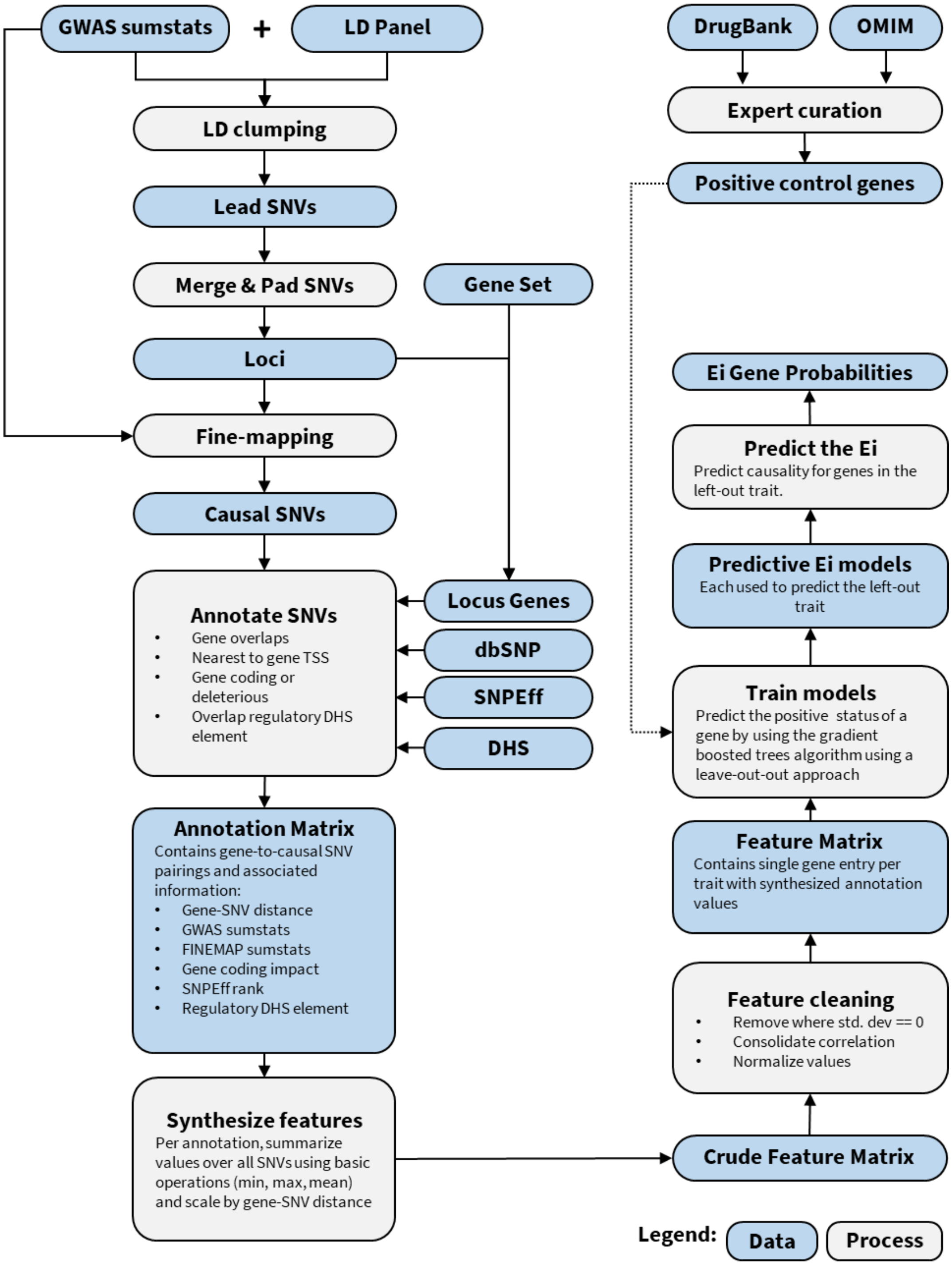


**Supplementary Fig. 9. Ei data and process workflow.** Input includes GWAS summary statistics and am appropriate LD reference panel. First, GWAS summary statistics are LD-clumped to determine a set of GWAS loci for statistical fine-mapping to determine a set of putatively causal SNVs at each locus. Genes overlapping the GWAS loci are paired to causal SNVs via their proximity to genes and further annotated as to whether the SNVs confer functional impact (using SNPEff, dbSNP for coding impact and DHS overlap for non-coding). Further information is added to the SNV-gene pairing such as gene length, GWAS summary statistics (eg. effectsize, MAF) and FINEMAP summary statistics (eg. log10(Bayes Factor)). The output consists of an annotation matrix where each row is an SNV-gene pairing and columns contain information regarding the SNV or gene, including SNV-gene distance (see **Supplemental Table 6**). The annotation matrix (see **Supplementary Table 6**) may contain multiple SNVs paired to a given gene and thus multiple annotation values (such as coding impact, GWAS effect size, etc) may be associated to each gene. To resolve this into a set of features, each annotation is summarized over all SNVs paired to a gene using basic operations (sum, min, mean, max) as well as scaling of these values based on their distance to the gene. A listing of features is provided in **Supplementary Table 7**. Model training was conducted using a leave-one-out approach, whereby each trait-specific model was trained using data from the remaining traits.
